# Blood Concentrations of Per- and Polyfluoroalkyl Substances are Associated with Autoimmunity-like Effects in the American Alligator

**DOI:** 10.1101/2022.02.15.480575

**Authors:** T. C. Guillette, Thomas W. Jackson, Matthew P. Guillette, James P. McCord, Scott M. Belcher

## Abstract

Surface and ground water of the Cape Fear River basin in central and coastal North Carolina is contaminated with high levels of per- and polyfluoroalkyl substances (PFAS). Elevated levels of PFAS have also been found in blood of fish and wildlife living in and around the Cape Fear River, and in the blood of human populations reliant on contaminated well or surface water from the Cape Fear River basin as a sources of drinking water. While the public and environmental health impacts of long-term PFAS exposures are poorly understood, elevated blood concentrations of some PFAS are linked with immunotoxicity and increased incidence of some chronic autoimmune diseases in human populations. The goal of this study was to evaluate PFAS exposure and biomarkers related to immune health in populations of American alligators (*Alligator mississippiensis*), a protected and predictive sentinel species of adverse effects caused by persistent toxic pollutants. We found that serum PFAS concentrations in a representative population of alligators from the Cape Fear River were increased compared to a reference population of alligators from the adjoining Lumber River basin. The elevated serum PFAS concentrations in the Cape Fear River alligators were associated with increased innate immune activities, and autoimmune-like phenotypes were observed in this population. In addition to evidence of significantly higher double stranded-DNA binding autoantibodies in adult Cape Fear River alligators, our qRT-PCR analysis found remarkably high induction of Interferon-α αsignature genes implicated in the pathology of human autoimmune disease. We interpret the association of increased PFAS exposure with disrupted immune functions to indicate that PFAS broadly alters immune activities resulting in autoimmune-like pathology in American alligators. This work substantiates and extends evidence from experimental models and human epidemiology studies showing that some PFAS are immune toxicants.

**Highlights:**

- Serum PFAS concentration were elevated in alligator populations from the Cape Fear River
- Higher PFAS concentrations were associated with disrupted innate immune functions
- *Infa*-responsive autoimmunity signature genes were induced in Cape Fear River alligators
- Elevated PFAS levels were associated with autoimmune-like activity in alligators 26

## 1. Introduction

Per- and polyfluoroalkyl substances (PFAS) are a class of synthetic organic chemicals that are global contaminants of both built and natural environments (OECD 2018). Owing to their extensively fluorinated aliphatic backbone, PFAS are chemically stable, resistant to thermal and enzymatic breakdown, and can persist in terrestrial and aquatic environments. Because of their widespread use over the past 70 years, many PFAS and their terminal breakdown products, have become ubiquitous contaminants of the land, water, and air through which humans and wildlife are exposed (De Silva et al., 2021). In the US alone, drinking water supplies for an estimated 200 million people are contaminated with PFAS, with recent analyses estimating the direct annual health-related cost due to PFAS exposure ranges from $37-59 billion in the US and €52-84 billion per year in the European Union (Andrews and Naidenko, 2020; Cordner et al., 2021) Evidence from experimental animal and human epidemiologic studies has linked increased exposure to the most abundant PFAS with immunotoxicity and altered immune functions (DeWitt, 2015). Those data led the United States National Toxicology Program to conclude that perfluorooctanoic acid (PFOA) and perfluorooctanesulfonic acid (PFOS) are hazards to the human immune system (DeWitt, 2015; NTP, 2016). Additional epidemiologic findings have also demonstrated associations between exposures to some PFAS with immunosuppression and adverse health impacts linked to increased incidence in childhood infection, decreased antibody production in response to vaccination, and increases in severity of COVID-19 (Fenton et al., 2021; Grandjean et al., 2020). There is also evidence from human studies that links some PFAS exposures with an increased incidence of chronic autoimmune disorders, including thyroid disease and inflammatory bowel diseases (Fenton et al., 2021). The mechanisms of PFAS-mediated immunotoxicity and their roles in autoimmunity are poorly understood.

The Cape Fear River basin, located in central and coastal North Carolina encompasses over 9,300 sq. miles of waterways that service ∼5.2 million people in rural and urban communities, and is typical in regard to contaminant burden of many regions with high PFAS contamination of surface, ground, and drinking water (Fig. 1A). PFAS contamination in the Cape Fear originates primarily from fluorochemical production, manufacturing, wastewater treatment discharges, and the use of aqueous film forming foams (AFFF) as fire suppressants (McCord and Strynar, 2019). In 2017, surface water sampling from the Cape Fear River revealed high levels of per- and polyfluoroether acids (PFEA) and fluoropolymer manufacturing byproducts with diverse chemical structures originating from a fluorotelomer and fluoropolymer PFAS production facility active since the mid-1980’s (Hopkins et al., 2018; Sun et al., 2016). Both novel PFEAs such as HFPO-DA (GenX), Nafion byproducts, and a variety of other fluorochemicals from PFAS-based aqueous film forming foam used as a fire suppressant by military, municipalities, and regional airports have now been detected in the drinking water and blood samples of residents living near the Cape Fear River (Kotlarz et al., 2020; McCord and Strynar, 2019; Ruyle et al., 2021). Recently, total organic fluorine analysis detected extremely high sum PFAS concentrations in excess of 100,000 ng/L in the Cape Fear River at drinking water treatment plant intakes suggesting that humans and aquatic ecosystems in the Cape Fear River basin have been experiencing very high levels of total PFAS exposure (Zhang et al., 2019). The potential for impacts on both ecosystems and human populations from this contamination has been confirmed by wildlife and human population based PFAS exposure studies that found increased levels of PFAS in fish, birds, and humans living near the Cape Fear River (Guillette et al 2020; Kotlarz et al. 2020; Robuck et al. 2021). The public and environmental health impacts of long-term exposures resulting from undocumented PFAS discharge are largely unknown.

**Fig 1.**
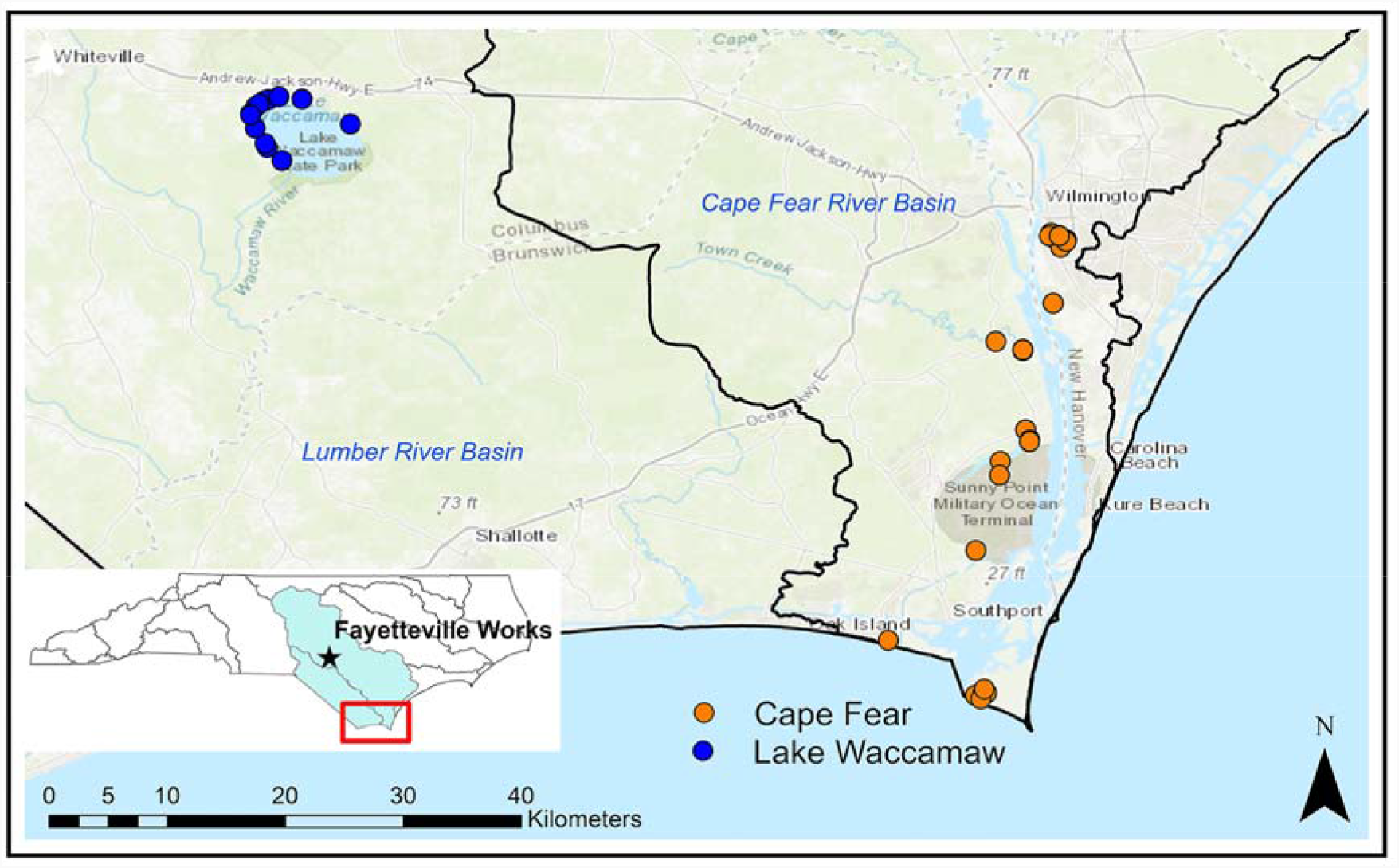
Location of study sites. Location of sampling sites collected at Lake Waccamaw (LW; blue) and from the Cape Fear River basin (CFR; orange). Blood samples and morphometric measures were collected from 26 alligators from LW and 49 from locations on the CFR. Some circles indicating sampling sites include multiple individuals. Inset shows the CFR basin in light blue with the location of known upstream fluorochemical production facility indicate with a star.

The aim of this study was to evaluate the impacts of long-duration PFAS exposure on biomarkers of immune health in American alligators (*Alligator* mississippiensis), a top trophic carnivore at the human/wildlife interface of both rural and urban environments (Beal and Rosenblatt, 2020; Somaweera et al., 2020). The American alligator has been used for more than four decades as an aquatic biomonitoring and predictive sentinel species of adverse outcomes resulting from the integrated effects of persistent toxic chemicals (Crain and Guillette 1998; Guillette et al. 2000; Pérez and Wise 2018). Alligators living along the Cape Fear River are non-migratory apex predators with a life span exceeding 60 years that co-utilize human habitats contaminated with PFAS (IUCN, 2018). Because innate and adaptive immune systems of American alligators are generally robust and highly protective (Finger and Gogal, 2013; Merchant and Britton, 2006), we hypothesized that populations of American alligators living in locations with elevated PFAS water contamination would be sensitive sentinels of cumulative adverse immune health effects resulting from PFAS exposures common to humans living in the Cape Fear River basin.

Building upon previous biomonitoring studies demonstrating utility of American alligators as effective indicators of environmental PFAS contamination (Bangma, et al 2017a; 2017b), the objective of the following study was twofold: 1) characterize the PFAS serum concentration profiles of two populations of alligators living in different watersheds suspected to have different PFAS contamination profiles and 2) examine associations between PFAS concentrations and immune health endpoints in alligators from these two watersheds. Given that NTP has classified two PFAS (PFOS and PFOA) as hazards to the immune system, we proposed that alligators would be a robust sentinel species to examine PFAS and immune endpoints, as American alligators have a robust and protective antibacterial and antiviral innate immune system (Merchant and Britton 2006; Finger and Gogal 2013). Previous research into the CFR aquatic ecosystem indicated a positive association between serum lysozyme, an important marker of the innate immune system, and serum PFOS concentrations in another aquatic predator, the Striped bass (Guillette el al 2020). Due to this finding, we expanded the immune endpoints examined in this study to include biomarkers of the adaptive immune system function, such as white blood cell counts and differentials, the presence of double-stranded DNA (dsDNA) autoantibodies, and Ifn-α responsive gene expression.

## 2. Methods

### 2.1 Animal and sampling procedures

All animal procedures were performed with approval by the North Carolina State University Institutional Animal Care and Use Committee (protocol #18-085-O). Alligators were sampled using active capture methods that employed snatch hooks from locations with direct access to the Cape Fear River and a site from the adjacent Lumber River basin, Lake Waccamaw, NC 134 (34.28726947673369, -78.50965117663293; Fig. 1). Cape Fear River sites included: Greenfield 135 Lake (34.2107388211135, -77.93707241283391) Wilmington, NC; Orton Pond in Brunswick 136 County (34.045198995423966, -77.98399224982941); Oak Island, NC (33.91736426475394, 137 -78.12476238375928), and Baldhead Island, NC (33.85295162890547, -77.97713359942439). Sampling occurred in 2018 (July 24 to October 25) and 2019 (April 18 to October 14).

Immediately following capture a 10-15 mL whole blood sample was collected from the post-occipital spinal venous sinus using a sterile 20 g or 18 g needle and a 30 mL syringe (Myburgh et al., 2014). Whole blood samples were transferred to 8 mL serum and lithium heparin coated plasma tubes (Vacutainer, BD, Franklin Lakes, NJ). Serum tubes were incubated for 30 min at ambient temperature to allow clot formation and then stored on ice. Plasma tubes were inverted gently following collection, then stored on ice until centrifugation. After field collections, blood was centrifuged (1800 x g for 10 min at 4°C) and serum/plasma fractions were immediately aliquoted into Teflon-free cryovials and stored at -80°C until analysis. Following the blood sample collections, alligators were visually examined for general health, and external injuries were noted and photographed. Sex was determined by cloacal examination with total length, snout to vent length (SVL), tail girth, head/snout, and rear foot measurements recorded. Following sample/data collection, alligators were released at the site of capture. Typical time from first contact to release of animal was dependent on animal size with the average time to release being < 15 min.

### 2.2. Chemical and Reagentss

Methanol (Optima^®^, Lot 183859), ammonium formate (99%, AC401152500), acetonitrile (ACN; Optima^®^, Lot 184819), and formic acid (99.5%, A117-50) from Fisher Scientific (Waltham, MA) was used for extractions and instrument solvent gradient. All water used for extraction, in aqueous buffers, and solutions were prepared in sterile Milli-Q A10 water (18Ω; 3 ppb total oxidizable organics), with the purified water analyzed for PFAS contamination prior to use. All laboratory glassware was rinsed with sterile Milli-Q water and methanol prior to use. The control material for the instrument methods was National Institute of Standards and Technology (NIST) Standard Reference Material (SRM) 1957 organic contaminants in non-fortified human serum. Chemicals for buffers and other standard laboratory chemicals were also purchased from Fisher Scientific and as indicated below.

### 2.3 LC-MS analysis of Serum PFAS Concentrations

Matrix matched (serum) calibration solutions (n = 23; ranging from 0.1 ng/mL to 100 ng/mL) were prepared from neat standards in charcoal stripped fetal bovine serum (Life Technologies, Grand Island, NY; cat #10437, Lot #1754113; total protein 3.7 g/dL). An internal standard (IS) solution was prepared by diluting a solution containing 10 mass-labeled PFAS in water with 0.1 M formic acid (99.5%, A117-50, Fisher Scientific, Waltham, MA): ^18^O_2_-PFBS,^13^C_4_-PFBA, ^18^O_2_-PFHxS, ^13^C_4_-PFOA, ^13^C_4_-PFOS, ^13^C_5_-PFNA, ^13^C_9_-PFDA, ^13^C_4_-PFBA, ^13^C_2_-PFHxA, ^13^C_2_-6:2FTS (Wellington Labs, Guelph, ON). An additional matrix matched quality control (QC) sample was made by combining randomly selected aliquots of sampled American alligator serum (n = 12). The resulting alligator QC sample and NIST SRM 1957 (n = 6) was used to test the method for reproducibility and stability (Table S2). Charcoal stripped fetal bovine serum was also spiked with 5 ng/mL of PFAS standards and used as an internal control for accurate measurement (n = 4). Additional control samples included fetal bovine serum method blanks (n = 7), and purified water field blanks (n = 5) made by injecting 4 mL of Milli-Q water into vacutainer tubes during field sampling on 5 different days during the sampling period.

All control and experimental samples were extracted and analyzed using methods identical to those detailed in our previous studies (Guillette et al. 2020). Briefly, 50 μl of each sample was aliquoted into a 2 mL polypropylene tube to which 100 μl of 0.1 M formic acid and internal standards (12.5 ng) were added. Ice-cold ACN (450 μl) was added to the tube and then vortexed for 3 sec. Samples were then centrifuged at 12,500 x g for 5 min at room temperature. The resulting supernatant (100 μl) was collected and added to 300 μl of aqueous 0.4 mmol ammonium formate and then transferred to polypropylene autosampler vials. PFAS were analyzed using a Vanquish UPLC system (Thermo Fisher Scientific, Waltham, MA) equipped with an Accucore C18+ column (2.1 mm x 100 mm x 1.5 μ) at a flow rate of 300 μL/min, injection volumes of 100 μL, and a binary mobile phase gradient composed of 95:5 H_2_O:ACN, 0.4 mM ammonium acetate and 95:5 ACN:H_2_O, 0.4 mM ammonium acetate. A Thermo Orbitrap Fusion mass spectrometer (Thermo Fisher Scientific, Waltham, MA) with a heated electrospray ionization (HESI) source operated in negative mode was used to detect PFAS. For compound validation, data was collected in data dependent mode with a preferred ion list consisting of the quantitated PFAS standards (Table S1). PFAS quantitation was based on an eight-point calibration curve with three injections per concentration randomly assigned in the sample run of the internal standard normalized integrated peak area of the extracted ion chromatogram of the [M-H]-ion with a 5-ppm mass tolerance. The r^2^ of all calibration curves used for analysis were above .97; limits of detection (LOD) were defined as the estimated mean concentration of method blanks plus three standard deviations (Table S2). Multiple replicates of SRM 1957 (n =6) were compared to the values on the Certificate of Analysis for the NIST SRM 1957 standards and were within 15.89 % of expected values (Table S4).

### 2.4 Lysozyme assay

The EnzChek® Lysozyme Assay (Thermo Fisher Scientific, Waltham, MA, Cat. E-22013) was used according to manufacturer’s protocols to determine lysozyme activity in serum of a subset of American alligator serum samples (Lake Waccamaw: n = 22, Wilmington CFR: n =21). Samples were diluted 1:5 in sample buffer and analyzed in triplicate, the average relative standard deviation (RSD) for QC and experimental samples were 7% and 9%, respectively.

### 2.5 Alligator serum/plasma complement assay

Complement activity was assessed using a modified sheep red blood cell lysis assay previously used in alligators to assess relative complement activity (Merchant and Britton 2006). Packed Sheep red blood cells (RBCs, Innovative Research Novi, MI) were aliquoted into 15 ml polypropylene conical tubes and centrifuged for 5 min at 800 x g. Cell pellets were washed with phosphate buffered saline (pH 7.4; PBS) and then gently suspend in PBS and recentrifuged. Washed cells were resuspended at a final concentration of 2% RBCs in PBS or PBS supplemented with a final concentration of 50 mM EDTA. Cells were rocked gently and 150 μl of the cell suspension was rapidly aliquoted into 1.7 ml microcentrifuge tubes containing 150 μl of each alligator serum sample and mixed by inversion. Control experiments with plasma samples were analyzed in parallel and yielded comparable results. Because sufficient volumes of plasma were not available for as many sampled animals, serum was used for the analysis. All sample aliquoting for each analysis was completed in ≤ 2 min. Mixed samples were incubated at room temperature for 20 min and then rapidly placed onto ice/water bath for 10 min, samples were centrifuged at 3000 x g for 5 min and then rapidly returned to ice. Supernatant (100 μl) was aliquoted in duplicate into clear 96-well microtiter plates. Absorbance at 540 nm was determined using a Tecan microplate reader. (Tecan Systems Inc., San Jose, CA) Controls on each analysis plate included a blank control containing 1x PBS only, 1% RBC in PBS, 1% RBC in 1x PBS supplemented with 50 mM EDTA, and a positive control of maximum RBC lysis containing 2% RBCs to which an equal volume of deionized water was added. Percent maximal complement activity for each sample was calculated by subtracting the mean absorbance from the EDTA containing samples from the mean absorbance of the sample lacking EDTA. That difference was divided by mean maximal absorbance of the positive control samples, and the product was multiplied by 100. Each sample was analyzed a minimum of two times. The mean % CV ± SD for maximal RBC lysis positive controls was 1.75 ± 1.42 % and for all experimental samples was 233 3.41 ± 4.43 %.

### 2.6 dsDNA ELISA analysis

Individual wells of 96 well microtiter plates were coated overnight at 4°C with 500 ng of double stranded calf thymus DNA suspended in DNA coating solution (Pierce, Rockford IL, cat # 17250). Plates were washed extensively with PBS containing 0.05 % Tween 20 (Fisher Scientific), incubated at room temperature for 2 hours in blocking buffer consisting of PBS supplemented with 10% bovine serum albumin (BSA; fraction V, Fisher Scientific), and then rewashed with PBS containing 0.05% Tween 20. Alligator plasma samples were diluted 1:500 and 1:1000 into PBS containing 10% BSA. Diluted samples (100 μl) were added to wells in duplicate or triplicate and incubated overnight at 4° C. Wells were then washed extensively with PBS/0.05% Tween 20, blotted dry by inversion on to paper towels, and then incubated for 1 hour at room temperature with affinity purified polyclonal horse radish peroxidase (HRP) conjugated goat anti-alligator IgG antiserum (Bethyl Labs, Montgomery, TX; Cat # A140-119P; Lot# A140-119P-6) that was diluted 1:10,000 (0.1 μg/ml) in PBS 10% BSA. Samples were extensively washed with PBS 0.05 % Tween 20, blotted dry, and 50 μl of 1x TMB Solution (Invitrogen, Cat # 00-4201-56) was added to each well. The peroxidase reaction was terminated following incubation for 30 min at room temperature by the addition of an equal 50 μl volume of 2M sulfuric acid, and the absorbance of each well measured at 450 nm. Control samples on each analyzed plate included calf serum diluted 1:500 (Gibco), blocking buffer only, 1:500 dilution of alligator plasma or calf serum sample lacking HRP-conjugated anti-alligator antiserum, and coated wells lacking DNA. Following optimization of alligator plasma concentrations and alligator antiserum dilutions, specificity of DNA binding was confirmed by observing significant decreases in immunoreactivity in DNA-coated wells that were treated with DNase1 (10 units; RQ1 Dnase, Promega Madison, WI), and in samples of alligator plasma pretreated with 100 μg of calf thymus DNA. Experimental plasma samples were analyzed in duplicate or triplicate on a single plate at 1:500 dilution and on the same plate using an independent dilution of 1:1000, with each analysis independently replicated. Representative samples selected randomly from samples with sufficient volume and quality of plasma from 4 females and 4 males from CFR and LW were analyzed. The average intra-assay coefficient of variation ± SD for all experimental samples was 5.9 ± 1.4% (1:500) and 4.5% ± 1.1% (1:1000). The average between plate coefficient of variation ± SD was 7.0 ± 1.7% (1:500) and 6.0 ± 1.5% (1:1000). The average absorbance 450 nM ± SD observed for each sample diluted at 1:500 was 2.05 ± 0.02 times greater than samples diluted at 1:1000.

### 2.7 Whole blood cell counts and leukocyte differential

Blood smears were prepared from ∼10 μl of peripheral whole blood taken from a subset of alligators. Blood smears were prepared in the field and allowed to air dry at room temperature on Superfrost plus slides (Fisher Scientific, Pittsburgh, PA). Slides were stained with Hemacolor® according to the manufacturer’s protocols (Sigma-Aldrich, St. Louis, MO; Cat: 1.11661). Peripheral whole blood cell counts were made by evaluation of stained blood smears using 40X and 100X objectives on a Nikon Eclipse 80i microscope (Nikon; Melville, NY). Each slide was examined independently by three different investigators blinded to sampling site, size, and sex. Leukocyte identity was determined by cellular morphology and staining characteristics (Sykes and Klaphake 2015; Stacy,et al., 2011). A minimum of 400 leukocytes and thrombocytes were counted to the calculate percentage of each cell type. Leukocytes were categorized into lymphocytes, basophils, heterophils, eosinophils, azurophils, and basophils with percent of each cell type determined by dividing the number of each cell type by the total cells counted minus the number of thrombocytes and then multiplying by 100. Variation between individual researcher counts was determined to be less than 7.9% for each cell type.

### 2.8 TBARS assay

Plasma lipid peroxidation was analyzed for 100 μl alligator plasma samples ran in duplicate using TBARS (TCA method) Assay kit (# 700870, Cayman Chemical, Ann Arbor, MI) according to manufacturer’s supplied protocols. Malondiadehyde concentrations were calculated by interpolation from a standard curve generated by serial dilution of manufacturer supplied standards.

### 2.9 Quantitative RT-PCR analysis

Total RNA from archived (stored at -80°C) American alligator whole blood was isolated using the Quick-RNA Whole Blood Kit (Zymo Research, Irvine, CA). Samples from both LW and CFR were selected randomly based only on the availability of samples. One µg of RNA was reverse transcribed using the high-capacity cDNA reverse transcription kit following manufacturer’s recommendations (Applied Biosystems; Grand Island, NY). Standard Fast TaqMan PCR amplification was performed in triplicate on a Step One Plus Real-Time PCR System (Applied Biosystems; Grand Island, NY) in a final volume of 20 µL containing ∼10 ng of cDNA (1.5 µL of RT product), 1x Universal Master Mix and custom TaqMan expression assay primers specific for each target alligator mRNA (**Table S5, S6**) (Applied Biosystems; Grand Island, NY). Relative expression was quantified using the 2^ΔΔCt^ method, in which ΔΔCt is the normalized value. Alligator *Gapdh* expression was used as an independent reference gene for normalization.

### 2.10 Data and statistical analysis

Each animal/blood sample was assigned a randomized numeric code by NC State University researchers at the time of collection. Samples were decoded for site and length/size/sex only after PFAS measurement/analysis and biological markers analysis were completed. Blood slides were further deidentified to mask sample identity for whole blood cell counts. Samples used for each analysis were randomly selected based only on availability of sufficient material for analysis, and when possible to balance analysis for comparable numbers of males and females, life stage (adult SVL > 90 cm; juvenile SVL < 90 cm), and to control for possible effects on seasonality samples were date matched.

Data was analyzed and visualized using Prism (version 9.3, GraphPad La Jolla, CA); R statistical programming environment version 3.5.2; SPSS (Version 27.0, IBM, Armonk, NY); JMP Genomics 9 (SAS, Cary, NC). For PFAS concentration data, means and concentration ranges were determined from values above the LOD and interpolated 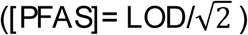 values were used when detected PFAS congeners were detectable but below the LOD (LOD = mean_blanks_ + 3(SD_blanks_), non-detects were replaced with a value of zero. A Shapiro-Wilk’s test was used to assess normality of data; for data requiring normalization a natural log transformation was used, [PFAS] data was log_10_ transformed, and percentage/ratiometric data were arcsine transformed. Non-parametric statistics were used for data not meeting analysis model assumptions following data transformation. Multivariate general linear modeling was used to examine PFAS concentrations with site and sex as fixed factors and SVL, month, and year as covariates. Linear regression and Pearson’s correlation coefficient or Spearman’s rank correlation coefficient were used to calculate relationships between total PFAS serum concentrations and serum biomarkers. A 2-way analysis of variance (ANOVA) was used to evaluate peripheral blood counts data (cell type, location), and ∑log_10_ [PFAS] (SVL, location) data and biomarker activities were initially evaluated at each site using multivariate ANOVA (biomarker activity, ∑ log_10_[PFAS], and/or location). Principal component analysis (PCA) was performed to dimensionally reduce PFAS concentrations using a restricted maximum likelihood estimation (REML) and components with a minimum eigenvalue of 1. Ward’s general agglomerative hierarchical clustering procedure was performed on serum PFAS congener concentrations (log_10_ transformed) to demonstrate large scale differences in PFAS clustering using JMP Genomics 9. A minimal level of statistical significance for differences in values among or between groups was considered p ≤ .05.

## 3. Results and Discussion

### 3.1 Analysis of serum PFAS concentrations

We used liquid chromatography and high-resolution mass spectrometry to determine concentrations of 23 different PFAS present in serum samples of 75 adult and juvenile alligators from North Carolina (Guillette et al. 2020). Alligators were sampled beginning in July 2018 and continued through October of 2019 at sites experiencing both point and non-point source PFAS exposures along the Cape Fear River (CFR; n = 49), and from Lake Waccamaw (LW; n = 26), a site < 50 km away and in the adjoining Lumber River watershed with no known fluorochemical production (Fig. 1). From the targeted list of 23 PFAS analyzed (Table S1), we found fourteen different PFAS, including long and short chain perfluoroalkyl acids (PFAA), perfluoroether acids (PFEA) and the fluorotelomer 6:2 FTS in these alligator blood samples (Table S2, S3).

As anticipated from previous wildlife exposure studies along the Cape Fear River, PFOS was the dominant PFAS detected in alligator serum (Guillette et al. 2020; Robuck et al. 2021) We detected PFOS in 100% of samples analyzed and it accounted for 79.7% and 75.8% of the total PFAS present in alligator serum samples from the CFR and LW respectively. The median number of PFAS congeners detected in serum samples from CFR alligators was 10 (range 4-12), whereas a median of 5 (range 2-9) were detected in LW samples (Table S3). The relative composition of long and short chain PFAAs and PFEAs detected in blood of CFR American alligators were in general agreement with those found in the blood of adults and children exposed to PFAS from drinking water in Wilmington, NC (Kotlarz et al., 2020). Results of our principal component analysis (Fig. 2), unsupervised hierarchal clustering (Fig.3A, 3B), and statistical analysis of total serum concentrations (Fig.3C) demonstrated both increases in PFAA concentrations and an exposure profile characteristic and predictive of samples from the CFR (Fig. 3A-C). Results of our general linear model demonstrated a significant overall effect of location (F = 3.2, p = .003) on PFAS concentrations. No effects were detected of SVL or sex on any PFAS concentration measured (Table S7). In the overall model, there was a significant difference in concentrations of PFBA (F = 2.6, p = .04), PFO4DA, (F = 3.9, p = .005), PFHxS (F = 3.3, p = .01), Nafion_bp2 (F = 4.3, p = .003), PFO5DoDA (F = 4.4, p = .003), PFNA (F = 4.6, p = .004), and PFDA (F = 3.0, p = .02). Although the differences in NVHOS (F = 2.0, p = .10), PFOS (F = 2.1, p = .08), or PFOA (F = 1.7, p = .16) concentration in the overall model was not statistically significant following correction (F = 2.1, p = .08), there was an effect of location on NVHOS (F = 7.0, p = .01), PFOS (F = 5.8, p = .02), and PFOA (F = 5.3, p = .03) concentration (Table S8). These results are consistent with a profile enriched for PFAS associated with chemical manufacturing facilities along the CFR. Concentrations and detection frequencies of PFEAs (HFPO-DA, Nafion byproduct-2), PFAS congeners related to upstream production and discharge from the Chemours Fayetteville Works facility, were exclusive to, or greatly enriched in the CFR serum samples (Fig. 3A, B; Table S2). By contrast the PFAS exposure profiles observed in blood of LW alligators were characterized by the presence of bioaccumulative six-carbon and longer PFAAs (Fig.3B; Table S8). However, the PFEAs PMPA (4, 15%), PDO4DA (6, 23%), and PFO5DoDa (5, 19%) were detectable in a some alligator serum samples from LW suggesting that there are low levels of PFEA contamination within the Lumber River basin ecosystem (Fig. 3B, C, Table S2). Because of the distinctive differences between LW and CFR samples, physiological endpoints were examined by two-way ANOVA using site and sex as fixed factors. As expected from the GLMM, concentrations of sum PFAS detected in serum of CFR alligator were elevated compared to alligators sampled from LW (Fig. 1E), two-way ANOVA found a significant effect of location on total serum PFAS concertation (F(1, 71) = 27.3, *p* < .0001). We did not detect differences in total PFAS concentrations between juveniles (SVL < 90 cm) and adults (F(1, 71) = 0.0018, *p* = .965) at either site. There was not an interaction between groups (F(1, 71) = 0.989, p = .323). Results of a protected Fisher’s LSD indicated that serum concentrations of PFAS were significantly greater in both the juveniles (*p* = .006) and adult alligators (*p* <.0001) from the CFR (Fig. 3C).

**Fig 2.**
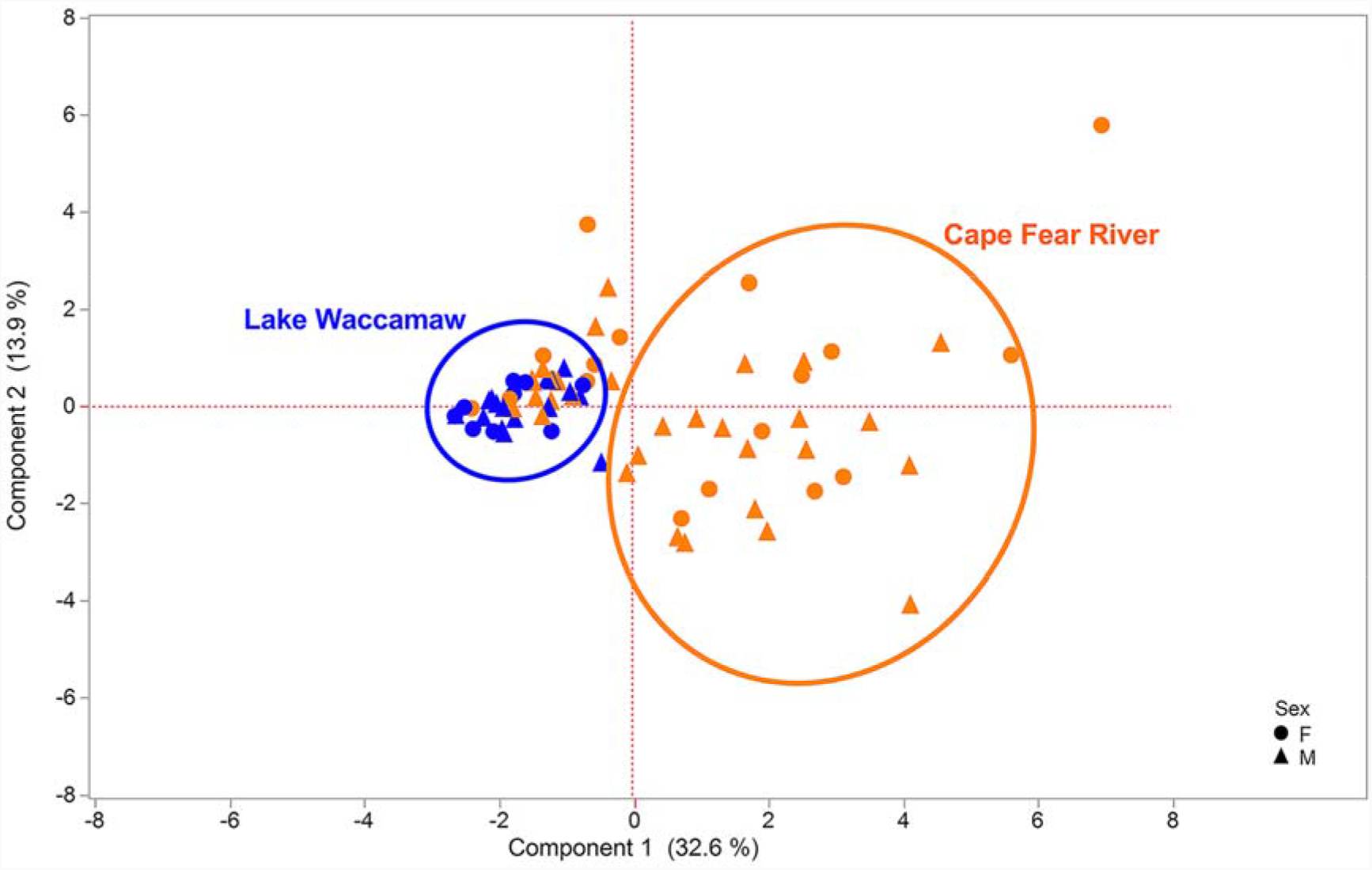
Principal component analysis. Principal component analysis (PCA) of log10 transformed serum PFAS congener concentrations from the CFR and LW was performed to dimensionally reduce PFAS concentrations using a restricted maximum likelihood estimation (REML) and components with a minimum eigenvalue of 1. Females are indicated with a circle and males are indicated with a triangle. Analysis used mean of log10 transformed PFAS concentrations (ng/ml) found in serum of adult (SVL ≥ 90 cm) and juvenile American alligators from LW (adult n = 15; juvenile n = 11) and the CFR (n = 26; juvenile = 23).

**Fig 3.**
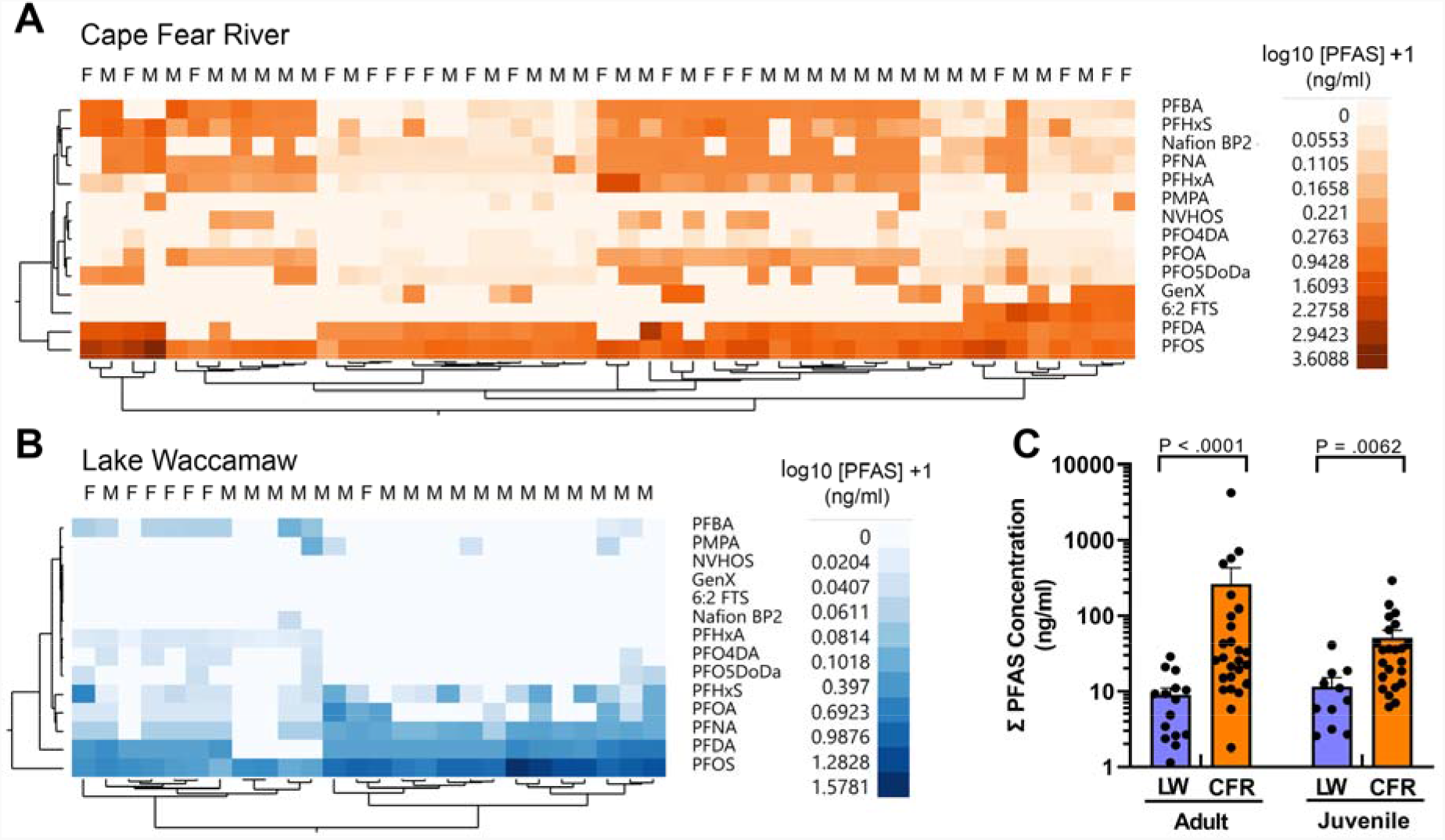
Serum PFAS exposure profiles. Shown are results of unsupervised hierarchal clustering of log10 transformed serum PFAS congener concentrations from (A) the CFR (orange) and (B) from LW (blue). Females are indicated with a capital F and males are indicated with a capital M. (C) Mean of PFAS concentrations (ng/ml) found in serum of adult (SVL ≥ 90 cm) and juvenile American alligators from LW (adult n = 15; juvenile n = 11) and the CFR (n = 26; juvenile = 23).

### 3.2 Associations between PFAS concentrations and innate immune activity

Initially focusing on lysozyme activity as a biomarker of innate immune function (Fig. 4A), the results of our two-way analysis of variance, with sampling site (CFR, LW) and size class (adult, juvenile) as between sample factors, revealed a main effect of age class (F (1, 44) = 13.35, *p* = .0007), the effect of sampling site was not significant (F (1, 44) = 0.14, *p* = .713). This was qualified by an interaction between sampling site and age class (F (1, 44) = 4.772, *p* = .03). A Fisher’s LSD multiple comparisons test indicated that lysozyme activity of LW juvenile alligators (*M* = 82.0 u/ml, SD = 20.7) was not significantly different than activity of CFR Juveniles (*M* = 70.7 u/ml, SD = 38.5; *p* = .24), however activity was significantly greater than activity found in adult alligators from both LW (*M* = 39.6 u/ml, SD = 17.3; *p* = .001) and the CFR (*M* = 57.4 u/ml, SD = 29.9; *p* = .02). Lysozyme activity of CFR adults was significantly higher than activity in adults from LW (Fig 4A; *p* = .05). Those findings were similar to our previous findings that higher concentrations of PFAS were associated with elevated lysozyme activity in striped bass (*Morone saxatilis*) from the CFR (Guillette et al. 2020). Using Pearson’s ranked correlation to assess the relationship between log10 transformed total PFAS concentrations and ln transformed lysozyme activity of adult alligators from both sites, we observed a positive correlation between serum PFAS and lysozyme activity, *r*(31) = .39, *p* = .006 (Fig. 4B). We did not observe a similar correlation between PFAS concentrations and lysozyme activity in juveniles (*r*(19) = -0.24.39, *p* = .334). While the differences in lysozyme activity in juveniles may be related shorter duration of PFAS exposure, ecological, population density, or other undefined factors including age or hormonal changes related to sexual maturity could be playing roles in these observed difference. Additional long-term biomonitoring and immune-health studies of will be required to elucidate the factors influencing the impacts of PFAS exposure on lysozyme activity.

**Figure 4.**
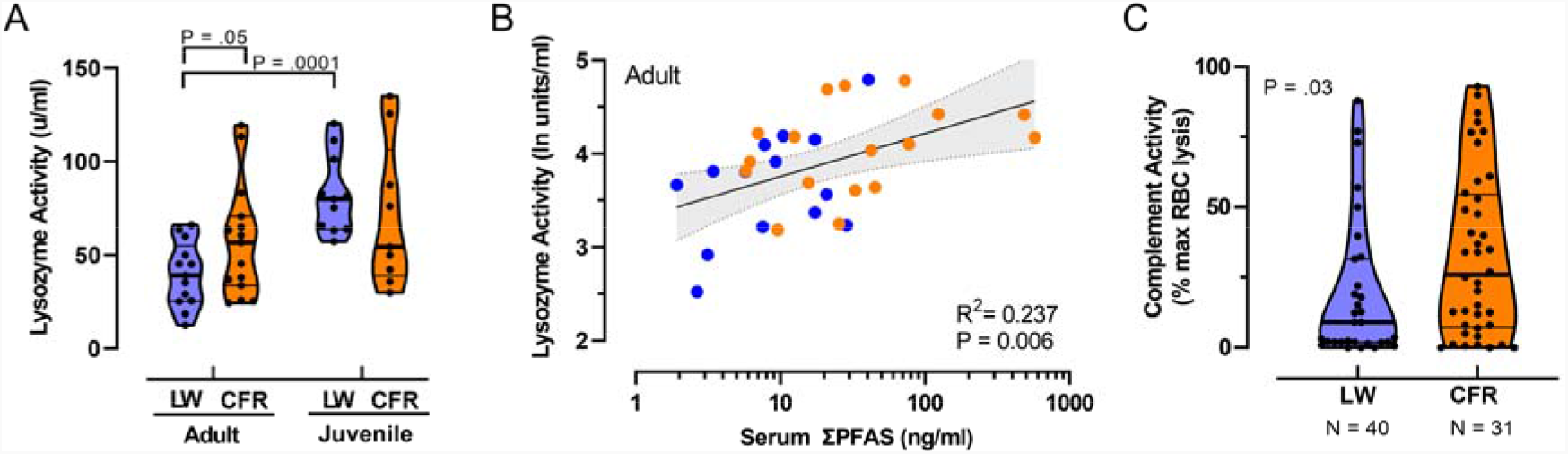
Impacts and relationship of PFAS exposure on innate immune functions. (A) Violin plots comparing median lysozyme activity in juvenile (SVL < 90 cm) and adult American alligators (SVL > 90 cm) from Lake Waccamaw (LW; thick lines indicate median; thin lines indicate quartiles) and from the Cape Fear River (CFR) LW adult n = 13, juvenile n = 11; CFR adult n = 15, juvenile n = 9. (B) Relationship between total PFAS concentrations and lysozyme activity in adult alligators. Solid line is the best fit linear regression, with shaded area represents 95% CI. n = 31 (C) Violin plots of complement activity in plasma samples of alligator from LW and CFR. Shown is the percent of maximal EDTA sensitive sheep red blood cell lysis activity (thick lines indicate median; thin lines indicate quartiles; LW: n = 40, CFR n = 31.

Using a Mann-Whitney test we found that relative serum complement activity, a key component of the humoral innate response linked with autoimmunity (Zipfel, 2009), was greater in CFR alligators (*Mdn* = 26.0) than for alligators sampled from LW (*Mdn* = 9.1), U = 436, *p* = .03 (Fig. 4C). As was observed for lysozyme, log10 transformed total PFAS concentrations and serum complement activity was moderately positively correlated, *r*(71) = .23, *p* = .05.

### 3.3 Adverse phenotypes observed in alligators from the CFR

Despite living in environments with near-constant exposure to pathogenic microorganisms, and having especially high body burdens of fecal coliform and other pathogenic bacteria, crocodilians, including wild American alligators, rarely suffer from systemic or skin infections from these microorganisms (Johnston et al., 2010; Keenan and Elsey, 2015; Zimmerman et al., 2010). However, during our sampling we observed a notably uncharacteristic increase in incidence of skin lesions, unhealed, and infected wounds in ∼20% of sampled CFR alligators (Fig. 5A). Field observations found purulent exudate and notable odor consistent with infection, and extensive slouth (white/gray devitalized tissue and debris) was frequently observed (Fig. 5A). The presence and appearance of these lesions were considered reminiscent of autoimmune disease related vasculitis in humans (Shanmugam et al., 2017). Although fresh and healed wounds, including loss of limbs that were likely related to territorial and mating interactions were observed in alligators sampled from LW, similarly infected and unhealed wounds were not observed. We also observed decreased clotting in some CFR alligators following blood collection (field observations); that phenotype was consistent with our findings of modestly decreased numbers of thrombocytes in whole blood cell counts from CFR alligators (Fig. 5B). A Mann-Whitney test indicated that that numbers of thrombocyte present in blood samples from LW were greater (*Mdn* = 27.25) than in samples from the CFR (*Mdn* = 22.30), *U* = 120.5, *p* =0.007.

**Fig. 5.**
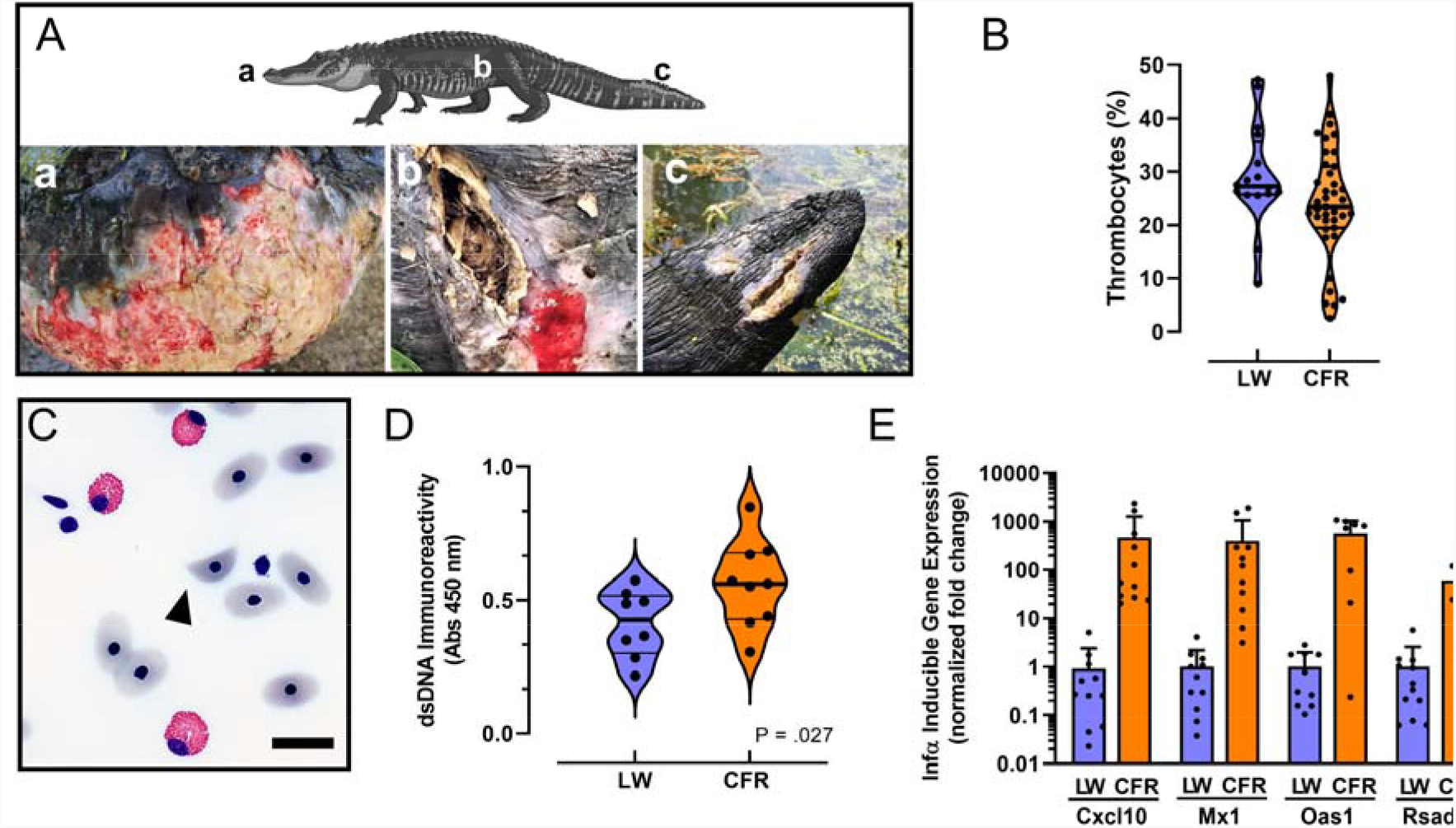
Autoimmune-like phenotypes associated with PFAS exposure. (A) Examples of cutaneous lesions and unhealed wounds associated with PFAS exposure in 3 representative animals sampled from the CFR in 2019. Lower case letters indicate body location of lesion shown in each panel. (B) Violin plots comparing median thrombocyte numbers quantified from whole blood smears in alligators from Lake Waccamaw (n = 16) and the Cape Fear River (n = 39). (C) Representative photomicrograph of a schistocyte (arrowhead) observed in a whole blood sample isolated from an alligator from the CFR. Bar = 10 μ m (D) Violin plots comparing median anti-dsDNA immunoreactivity in male (n = 4) and female (n = 4) alligator serum samples from each site. (E) Quantitative RT-PCR analysis of Infα signature gene expression analysis. Note the X-axis scale is Log10 relative fold-change. All differences in expression are significant (P < .0001). For violin plots median is indicted by a thick line and thin lines indicate quartiles.

### 3.4 Leukocyte and red blood cell phenotypes

Our analysis of differential white blood cell counts found no evidence for a general increase in chronic stress; a Mann-Whitney test found no significant differences in percent of heterophiles (*Mdn =* 16.7%) and LW (*Mdn* = 15.7%), *U* = 159, *p* = .09, and the heterophile/lymphocyte ratios from CFR (*Mdn* = 0.370) and LW (*Mdn* = 0.358) were also not significantly different, *U* = 194, *p* = .378. We also compared the relative levels of lipid peroxidation products in plasma and found no evidence for increases in general systemic mechanisms of cellular injury and stress in the CFR alligators (*Fig*. S1; LW *Mdn* = 1.60 μM, CFR *Mdn* = 1.51 μM; *U* = 359, *p* = .757).

During our analysis of white blood cells, we observed a greater number of samples with abnormal red blood cell phenotypes including schistocytes (helmet cells) in samples from CFR alligators. Helmet cells were found in 65.5% of CFR samples (n = 29, Fig. 5C), whereas none were found in the samples analyzed from LW (n = 9; Table S2). A Fisher’s exact test indicated that the observed number of samples from CFR alligators with schistocytes was significantly greater (*p* = .001) compared to the samples from LW. Schistocytes are fragmented red blood cell that in humans most often arise from mechanical shear forces caused by damaged endothelium and is a characteristic feature of microangiopathic hemolytic anemia. These abnormal blood cells are often associated with autoimmune disorders including lupus nephritis in humans (Anders et al., 2020; Tefferi and Elliott, 2004).

### 3.5 dsDNA binding antibodies and Infα-responsive autoimmune signature gene expression

Based on the observed presence of helmet cells and modestly lower numbers of thrombocytes associated with elevated PFAS exposure in CFR alligators, we evaluated other hallmark phenotypes associated with human autoimmune disorders (Crow et al., 2019; Sirobhushanam et al, 2021). Using an alligator specific anti-double stranded DNA (dsDNA) immunoglobulin (IgG) binding enzyme-linked immunosorbent assay, we evaluated a subset of male and female alligators from the CFR and LW and found that dsDNA binding IgGs were significantly greater in CFR alligator plasma samples (*M* = 0.577, *SD* = 0.167), *t*(14) = 2.22, *p* = .04, compared to LW samples (*M* = 0.413, *SD* = 0.127; Fig. 5D). Finally, our RT-PCR analysis of type 1 interferon (Ifn-α) responsive gene expression in alligator whole blood samples found extremely high expression of four different Ifn-α signature genes implicated in pathology of human autoimmune diseases including systemic lupus erythematosus (Crow et al, 2019). Relative to samples from LW, *Ifn-α* responsive gene expression was induced > 60-fold for *Rsad2* (viperin), and > 400-fold for *Cxcl10, Mx1*, and *Oas1* in CFR alligator blood samples (Fig. 5E; Table S5). The finding of extremely high induction of Inf-α signature gene expression in the CFR alligators supports a central role for excessive production of type I interferons in increasing the innate immune functions associated with elevated PFAS exposures. We interpret the greatly increased Inf-α signature gene expression, elevated autoantibodies, vasculitis-like skin lesions, elevated incidence of unhealed wounds, and elevated innate immune activity observed in the CFR population as indicative of elevated autoimmune-like activities in this population of American alligators.

## 4. Conclusions

Overall, our work found associations across multiple hallmark phenotypes of autoimmune related endpoints, some of which were shown to be associated with elevated PFAS concentrations in American alligators from the CFR. The PFAS-associated changes in both the innate immune and adaptive immune system may represent a significant health concern for wildlife and human health that may leave exposed populations at greater risk for infections and autoimmune disease. Uniquely, for this work we implemented an integrated One Environmental Health approach using American alligators as sentinels to more fully understand the consequences of long-term exposure to complex and poorly understood mixtures of PFAS with the aim of improving health of impacted human, animals, and ecosystems (Gibbs 2014; Pérez and Pierce 2018). In addition to being long lived, and sharing their environment with humans, American alligators and other crocodilians, by virtue of having innate immune functions optimized to eliminate microbial pathogens, may serve as sensitive and predictive model for detecting autoimmune hazards caused by chemical pollutants. Our findings of adverse immune-health effects in American alligators that have exposure profiles similar to the surrounding human population illuminates and reaffirms the need to reduce exposure and cease production and use of a chemical class that, through its ubiquity and persistence, is a global environmental health concern.

## Abbreviations

ACN: acetonitrile
CFR: Cape Fear River
CV: coefficient of variation
F: female; Infα interferon alpha type 1
IS: internal standard
LW: Lake Waccamaw
LOD: limit of detection
M: male:
PCA: principal component analysis
RBC: red blood cell
SLE: systemic lupus erythematosus
SRM: standard reference material
SVL: snout to vent length
TBARS: thiobarbituric acid reactive substances

## Acknowledgments

The authors would like to dedicate this manuscript to the memory of Dr. Louis J. Guillette Jr who continues to inspire us all.

We would also like to thank the staff of North Carolina Wildlife Resource Commission, especially Alicia Davis, Christopher Kent, and John Henry Harrelson. We are grateful for the help and tolerance we received from the residents of Wilmington, Lake Waccamaw, Bald Head Island, and Oak Island, NC. We are also indebted to our many volunteers, including Deborah and Stephen Burke, Kathy Sykes, Frank Robb, our community partners from the Bald Head Island Conservancy and Cape Fear River Watch who helped with this study, and acknowledge the effort of former lab members Gabe Bendfeldt, Helen Nguyen, Madi Polera, Aubrey L Sasser, and Chris Scheibly who enthusiastically contributed to sampling and data collection. We also thank Dr. Mark Stryner for his analytical assistance, and Dr. Heather Patisaul, Dr. Kylie Rock, and Hannah Starnes for critical review of the manuscript. Although EPA employees contributed to this article, the research presented was not funded by EPA and was not subject to EPA’s quality system requirements. Consequently, the views, interpretations, and conclusions expressed in the article are solely those of the authors and do not necessarily reflect or represent EPA’s views or policies.

## Funding

Research reported in this publication was supported by the National Institute Of Environmental Health Sciences of the National Institutes of Health under Award Number P42ES031009, P30 ES025128, and T32ES007046. The content is solely the responsibility of the authors and does not necessarily represent the official views of the National Institutes of Health. Additional support was funded by a NC Sea Grant Community Collaborative Research Grant (SMB), the NC Policy Collaboratory PFAS Testing Network (SMB).

## Author contributions

Conceptualization: SMB, TCG, MG

Methodology: SMB, TCG, MG, TJ, JM

Investigation: SMB, TCG, MG

Visualization: SMB, TCG, TJ

Funding acquisition: SMB

Project administration: SMB

Supervision: SMB

Writing – original draft: SMB, TCG

Writing – review & editing: SMB, TCG, MG, TJ, JM

## Data and materials availability

All data, code, and materials used in the analysis will be made available for purposes of reproducing or extending the analysis. All data are available in the main text and the supplementary materials.

## Notes

**Conflicts of Interest:** The authors declare they have nothing to disclose.

### Competing Interest Statement

The authors have declared no competing interest.

### Summary of Updates

Re-submission to a different journal. Changes based on reviewers comments

## References

Anders, H.-J., Saxena, R., Zhao, M.-H., Parodis, I., Salmon, J.E., Mohan, C., 2020. Lupus nephritis. Nat Rev Dis Primers 6, 7. https://doi.org/10.1038/s41572-019-0141-9

Andrews, D.Q., Naidenko, O.V., 2020. Population-Wide Exposure to Per- and Polyfluoroalkyl Substances from Drinking Water in the United States. Environ. Sci. Technol. Lett. 7, 931–936. https://doi.org/10.1021/acs.estlett.0c00713

Bangma, J.T., Bowden, J.A., Brunell, A.M., Christie, I., Finnell, B., Guillette, M.P., Jones, M., Lowers, R.H., Rainwater, T.R., Reiner, J.L., Wilkinson, P.M., Guillette, L.J., 2017a. Perfluorinated alkyl acids in plasma of American alligators (Alligator mississippiensis) from Florida and South Carolina. Environ Toxicol Chem 36, 917–925. https://doi.org/10.1002/etc.3600

Bangma, J.T., Reiner, J.L., Jones, M., Lowers, R.H., Nilsen, F., Rainwater, T.R., Somerville, S., Guillette, L.J., Bowden, J.A., 2017b. Variation in perfluoroalkyl acids in the American alligator (Alligator mississippiensis) at Merritt Island National Wildlife Refuge. Chemosphere 166, 72–79. https://doi.org/10.1016/j.chemosphere.2016.09.088

Beal, E.R., Rosenblatt, A.E., 2020. Alligators in the big city: spatial ecology of American alligators (Alligator mississippiensis) at multiple scales across an urban landscape. Sci Rep 10, 16575. https://doi.org/10.1038/s41598-020-73685-x

Cordner, A., Goldenman, G., Birnbaum, L.S., Brown, P., Miller, M.F., Mueller, R., Patton, S., Salvatore, D.H., Trasande, L., 2021. The True Cost of PFAS and the Benefits of Acting Now. Environ. Sci. Technol. 55, 9630–9633. https://doi.org/10.1021/acs.est.1c03565

Crain, D.A., Guillette, L.J., 1998. Reptiles as models of contaminant-induced endocrine disruption. Anim Reprod Sci 53, 77–86. https://doi.org/10.1016/s0378-4320(98)00128-6

Crow, M.K., Olferiev, M., Kirou, K.A., 2019. Type I Interferons in Autoimmune Disease. Annu Rev Pathol 14, 369–393. https://doi.org/10.1146/annurev-pathol-020117-043952

De Silva, A.O., Armitage, J.M., Bruton, T.A., Dassuncao, C., Heiger-Bernays, W., Hu, X.C., Kärrman, A., Kelly, B., Ng, C., Robuck, A., Sun, M., Webster, T.F., Sunderland, E.M., 2021. PFAS Exposure Pathways for Humans and Wildlife: A Synthesis of Current Knowledge and Key Gaps in Understanding. Environ Toxicol Chem 40, 631–657. https://doi.org/10.1002/etc.4935

DeWitt, J.C. (Ed.), 2015. Toxicological Effects of Perfluoroalkyl and Polyfluoroalkyl Substances, Molecular and Integrative Toxicology. Springer International Publishing, Cham. https://doi.org/10.1007/978-3-319-15518-0

Fenton, S.E., Ducatman, A., Boobis, A., DeWitt, J.C., Lau, C., Ng, C., Smith, J.S., Roberts, S.M., 2021. Per- and Polyfluoroalkyl Substance Toxicity and Human Health Review: Current State of Knowledge and Strategies for Informing Future Research. Environ Toxicol Chem 40, 606–630. https://doi.org/10.1002/etc.4890

Finger, J.W., Gogal, R.M., 2013. Endocrine-disrupting chemical exposure and the American alligator: a review of the potential role of environmental estrogens on the immune system of a top trophic carnivore. Arch Environ Contam Toxicol 65, 704–714. https://doi.org/10.1007/s00244-013-9953-x

Gibbs, E.P.J., 2014. The evolution of One Health: a decade of progress and challenges for the future. Veterinary Record 174, 85–91. https://doi.org/10.1136/vr.g143

Grandjean, P., Timmermann, C.A.G., Kruse, M., Nielsen, F., Vinholt, P.J., Boding, L., Heilmann, C., Mølbak, K., 2020. Severity of COVID-19 at elevated exposure to perfluorinated alkylates. PLoS One 15, e0244815. https://doi.org/10.1371/journal.pone.0244815

Guillette, L.J., Crain, D.A., Gunderson, M.P., Kools, S.A.E., Milnes, M.R., Orlando, E.F., Rooney, A.A., Woodward, A.R., 2000. Alligators and Endocrine Disrupting Contaminants: A Current Perspective. Am Zool 40, 438–452. https://doi.org/10.1093/icb/40.3.438

Guillette, T.C., McCord, J., Guillette, M., Polera, M.E., Rachels, K.T., Morgeson, C., Kotlarz, N., Knappe, D.R.U., Reading, B.J., Strynar, M., Belcher, S.M., 2020a. Elevated levels of per- and polyfluoroalkyl substances in Cape Fear River Striped Bass (Morone saxatilis) are associated with biomarkers of altered immune and liver function. Environ Int 136, 105358. https://doi.org/10.1016/j.envint.2019.105358

Guillette, T.C., McCord, J., Guillette, M., Polera, M.E., Rachels, K.T., Morgeson, C., Kotlarz, N., Knappe, D.R.U., Reading, B.J., Strynar, M., Belcher, S.M., 2020b. Elevated levels of per- and polyfluoroalkyl substances in Cape Fear River Striped Bass (Morone saxatilis) are associated with biomarkers of altered immune and liver function. Environ Int 136, 105358. https://doi.org/10.1016/j.envint.2019.105358

Hopkins, Z.R., Sun, M., DeWitt, J.C., Knappe, D.R.U., 2018. Recently Detected Drinking Water Contaminants: GenX and Other Per- and Polyfluoroalkyl Ether Acids: JOURNAL AWWA. Journal - American Water Works Association 110, 13–28. https://doi.org/10.1002/awwa.1073

IUCN, 2018. Alligator mississippiensis: Elsey, R., Woodward, A. & Balaguera-Reina, S.A.: The IUCN Red List of Threatened Species 2019: e.T46583A3009637. https://doi.org/10.2305/IUCN.UK.2019-2.RLTS.T46583A3009637.en

Johnston, M.A., Porter, D.E., Scott, G.I., Rhodes, W.E., Webster, L.F., 2010. Isolation of faecal coliform bacteria from the American alligator (Alligator mississippiensis). J Appl Microbiol 108, 965–973. https://doi.org/10.1111/j.1365-2672.2009.04498.x

Keenan, S.W., Elsey, R.M., 2015. The Good, the Bad, and the Unknown: Microbial Symbioses of the American Alligator. Integrative and Comparative Biology 55, 972–985. https://doi.org/10.1093/icb/icv006

Kotlarz, N., McCord, J., Collier, D., Lea, C.S., Strynar, M., Lindstrom, A.B., Wilkie, A.A., Islam, J.Y., Matney, K., Tarte, P., Polera, M.E., Burdette, K., DeWitt, J., May, K., Smart, R.C., Knappe, D.R.U., Hoppin, J.A., 2020. Measurement of Novel, Drinking Water-Associated PFAS in Blood from Adults and Children in Wilmington, North Carolina. Environ Health Perspect 128, 77005. https://doi.org/10.1289/EHP6837

McCord, J., Strynar, M., 2019. Identification of Per- and Polyfluoroalkyl Substances in the Cape Fear River by High Resolution Mass Spectrometry and Nontargeted Screening. Environ Sci Technol 53, 4717–4727. https://doi.org/10.1021/acs.est.8b06017

Merchant, M., Britton, A., 2006. Characterization of serum complement activity of saltwater (Crocodylus porosus) and freshwater (Crocodylus johnstoni) crocodiles. Comp Biochem Physiol A Mol Integr Physiol 143, 488–493. https://doi.org/10.1016/j.cbpa.2006.01.009

Myburgh, J.G., Kirberger, R.M., Steyl, J.C.A., Soley, J.T., Booyse, D.G., Huchzermeyer, F.W., Lowers, R.H., Guillette, L.J., 2014. The post-occipital spinal venous sinus of the Nile crocodile Crocodylus niloticus: its anatomy and use for blood sample collection and intravenous infusions. J S Afr Vet Assoc 85, e1–e10. https://doi.org/10.4102/jsava.v85i1.965

NTP, 2016. Monograph on Immunotoxicity Associated with Exposure to Perfluorooctanoic acid (PFOA) and perfluorooctane sulfonate (PFOS). National Toxicology Program.

OECD, 2018. New Comprehensive Global Database of Per- and Polyfluoroalkyl Substances.

Pérez, A., Pierce Wise Sr., J., 2018. One Environmental Health: an emerging perspective in toxicology. F1000Res 7, 918. https://doi.org/10.12688/f1000research.14233.1

Robuck, A.R., McCord, J.P., Strynar, M.J., Cantwell, M.G., Wiley, D.N., Lohmann, R., 2021. Tissue-specific distribution of legacy and novel per- and polyfluoroalkyl substances in juvenile seabirds. Environ Sci Technol Lett 8, 457–462. https://doi.org/10.1021/acs.estlett.1c00222

Ruyle, B.J., Thackray, C.P., McCord, J.P., Strynar, M.J., Mauge-Lewis, K.A., Fenton, S.E., Sunderland, E.M., 2021. Reconstructing the Composition of Per- and Polyfluoroalkyl Substances in Contemporary Aqueous Film-Forming Foams. Environ Sci Technol Lett 8, 59–65. https://doi.org/10.1021/acs.estlett.0c00798

Shanmugam, V.K., Angra, D., Rahimi, H., McNish, S., 2017. Vasculitic and autoimmune wounds. J Vasc Surg Venous Lymphat Disord 5, 280–292. https://doi.org/10.1016/j.jvsv.2016.09.006

Sirobhushanam, S., Lazar, S., Kahlenberg, J.M., 2021. Interferons in Systemic Lupus Erythematosus. Rheum Dis Clin North Am 47, 297–315. https://doi.org/10.1016/j.rdc.2021.04.001

Somaweera, R., Nifong, J., Rosenblatt, A., Brien, M.L., Combrink, X., Elsey, R.M., Grigg, G., Magnusson, W.E., Mazzotti, F.J., Pearcy, A., Platt, S.G., Shirley, M.H., Tellez, M., van der Ploeg, J., Webb, G., Whitaker, R., Webber, B.L., 2020. The ecological importance of crocodylians: towards evidence-based justification for their conservation. Biol Rev Camb Philos Soc 95, 936–959. https://doi.org/10.1111/brv.12594

Stacy, N.I., Alleman, A.R., Sayler, K.A., 2011. Diagnostic hematology of reptiles. Clin Lab Med 31, 87–108. https://doi.org/10.1016/j.cll.2010.10.006

Sun, M., Arevalo, E., Strynar, M., Lindstrom, A., Richardson, M., Kearns, B., Pickett, A., Smith, C., Knappe, D.R.U., 2016. Legacy and Emerging Perfluoroalkyl Substances Are Important Drinking Water Contaminants in the Cape Fear River Watershed of North Carolina. Environmental Science & Technology Letters 3, 415–419. https://doi.org/10.1021/acs.estlett.6b00398

Sykes, J.M., Klaphake, E., 2015. Reptile Hematology. Clin Lab Med 35, 661–680. https://doi.org/10.1016/j.cll.2015.05.014

Tefferi, A., Elliott, M.A., 2004. Schistocytes on the Peripheral Blood Smear. Mayo Clinic Proceedings 79, 809. https://doi.org/10.4065/79.6.809

Williams, A.J., Grulke, C.M., Edwards, J., McEachran, A.D., Mansouri, K., Baker, N.C., Patlewicz, G., Shah, I., Wambaugh, J.F., Judson, R.S., Richard, A.M., 2017. The CompTox Chemistry Dashboard: a community data resource for environmental chemistry. J Cheminform 9, 61. https://doi.org/10.1186/s13321-017-0247-6

Zhang, C., Hopkins, Z.R., McCord, J., Strynar, M.J., Knappe, D.R.U., 2019. Fate of Per- and Polyfluoroalkyl Ether Acids in the Total Oxidizable Precursor Assay and Implications for the Analysis of Impacted Water. Environ. Sci. Technol. Lett. 6, 662–668. https://doi.org/10.1021/acs.estlett.9b00525

Zimmerman, L.M., Vogel, L.A., Bowden, R.M., 2010. Understanding the vertebrate immune system: insights from the reptilian perspective. Journal of Experimental Biology 213, 661–671. https://doi.org/10.1242/jeb.038315

Zipfel, P.F., 2009. Complement and immune defense: From innate immunity to human diseases. Immunology Letters 126, 1–7. https://doi.org/10.1016/j.imlet.2009.07.005

