## supplemental methods for "Blood Concentrations of Per- and Polyfluoroalkyl Substances are Associated with Autoimmunity-like Effects in the American Alligator"

**This PDF file includes:**

Figs. S1

Tables S1 to S7

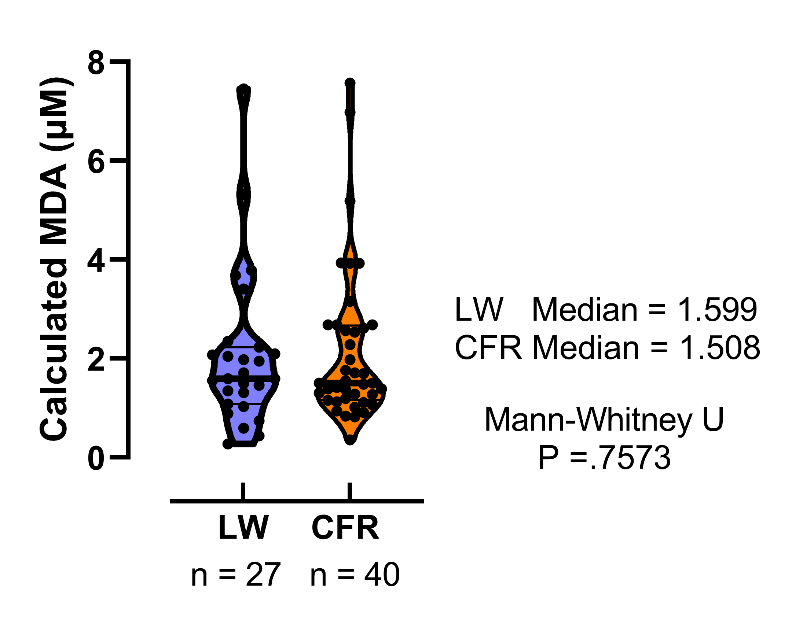

**Fig. S1.** Analysis of plasma lipid peroxidation: TBARS analysis.

**Table S1. Targeted List of 23 PFAS Analytes**

| Compound Name | Class | Family | Chemical Name | CAS # | Formula | Degree of Fluorination | Ether Bonds | Isomer Type | Functional Group | | Structure |
| --- | --- | --- | --- | --- | --- | --- | --- | --- | --- | --- | --- |
|  |  |  |  |  |  |  |  |  | R-CO_2_H^a^ | R-SO_3_H^b^ |  |
| PFBA | PFAA | PFCA | Perfluorobutanoic acid | 375-22-4 | C_4_HF_7_O_2_ | Per | 0 | Linear | Y | -- | 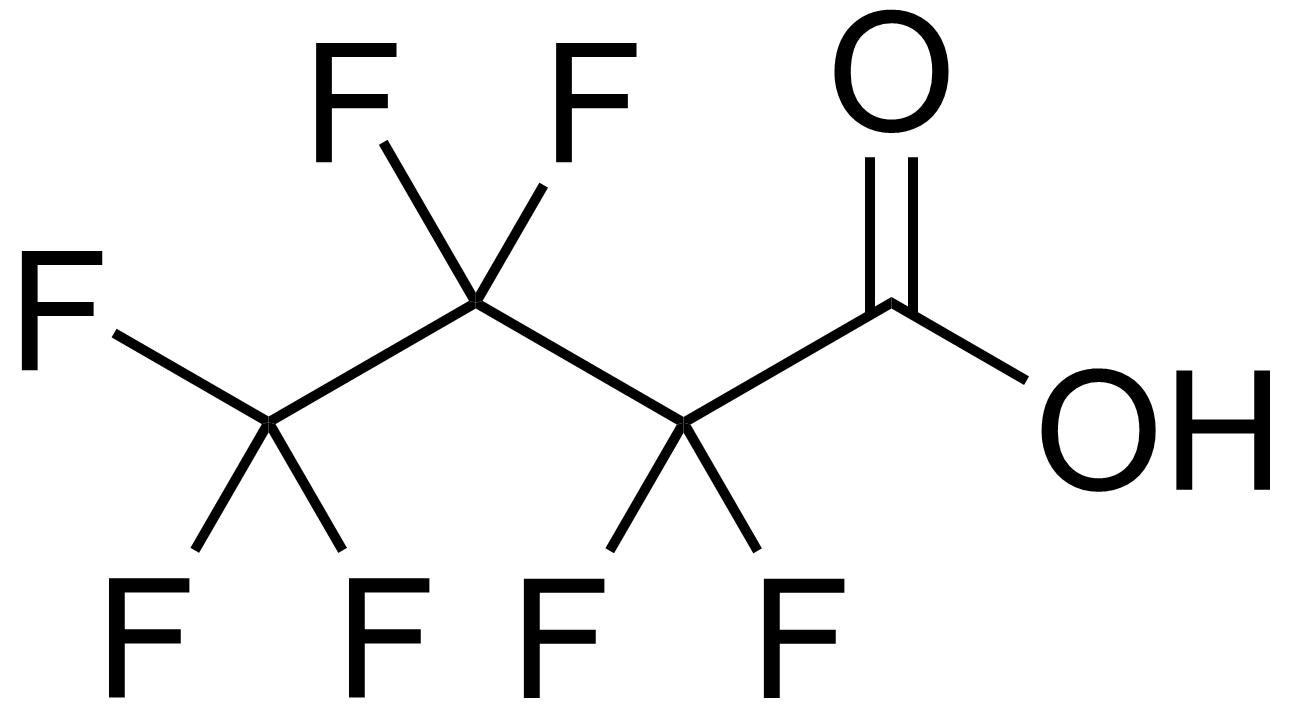 |
| PFPeA | PFAA | PFCA | Perfluoropentanoic acid | 2706-90-3 | C_5_HF_9_O_2_ | Per | 0 | Linear | Y | -- | 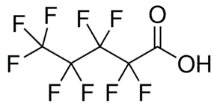 |
| PFHxA | PFAA | PFCA | Perfluorohexanoic acid | 307-24-4 | C_6_HF_11_O_2_ | Per | 0 | Linear | Y | -- | 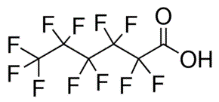 |
| PFHpA | PFAA | PFCA | Perfluoroheptanoic acid | 375‐85‐9 | C_7_HF_13_O_2_ | Per | 0 | Linear | Y | -- | 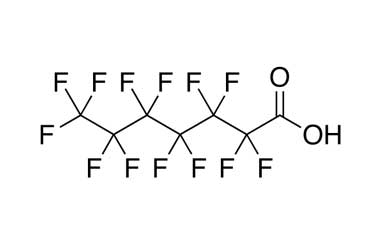 |
| PFOA | PFAA | PFCA | Perfluorooctanoic acid | 335-67-1 | C_8_HF_15_O_2_ | Per | 0 | Linear | Y | -- | 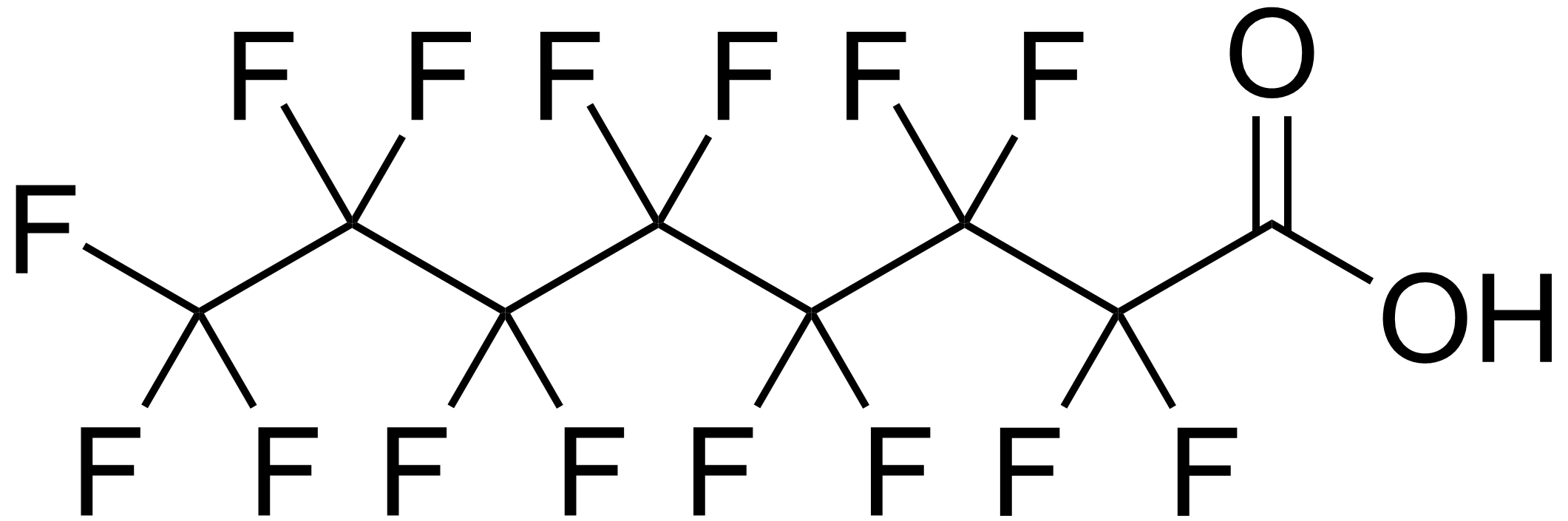 |
| PFNA | PFAA | PFCA | Perfluorononanoic acid | 375-95-1 | C_9_HF_17_O_2_ | Per | 0 | Linear | Y | -- | 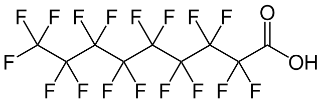 |
| PFDA | PFAA | PFCA | Perfluorodecanoic acid | 335-76-2 | C_10_HF_19_O_2_ | Per | 0 | Linear | Y | -- | 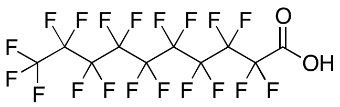 |
| PFBS | PFAA | PFSA | Perfluorobutanesulfonic acid | 375-73-5 | C_4_HF_9_SO_3_ | Per | 0 | Linear | -- | Y | 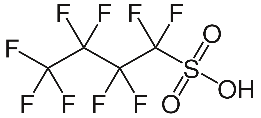 |
| PFHxS | PFAA | PFSA | Perfluorohexanesulfonic acid | 355-46-4 | C_6_HF_13_SO_3_ | Poly | 0 | Linear | -- | Y | 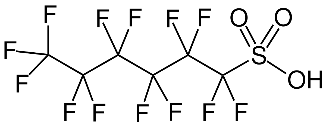 |
| PFOS | PFAA | PFSA | Perfluorooctanesulfonic acid | 1763-23-1 | C_8_HF_17_SO_3_ | Per | 0 | Linear | -- | Y | 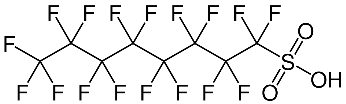 |
| PMPA* | PFAA | PFECA | Perfluoro-2-(perfluoromethoxy)propanoic acid | 13140-29-9 | C_4_HF_7_O_3_ | Per | 1 | Branched | Y | -- | 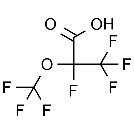 |
| PEPA* | PFAA | PFECA | Perfluoroethoxypropyl carboxylic acid | 267239-61-2 | C_5_HF_9_O_3_ | Per | 1 | Branched | Y | -- | 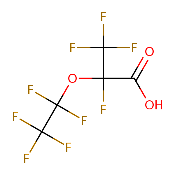 |
| HFPO-DA (GenX)* | PFAA | PFECA | Hexafluoropropylene oxide dimer acid | 13252-13-6 | C_6_HF_11_O_3_ | Per | 1 | Branched | Y | -- | 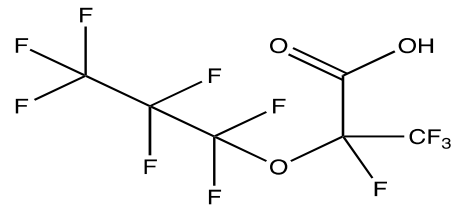 |
| PFO2HxA* | PFAA | PFECA | Perfluoro  (3,5-dioxahexanoic) acid | 39492-88-1 | C_4_HF_7_O_4_ | Per | 2 | Linear | Y | -- | 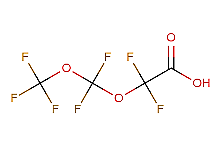 |
| PFO3OA* | PFAA | PFECA | Perfluoro  (3,5,7-trioxaoctanoic) acid | 39492-89-2 | C_5_HF_9_O_5_ | Per | 3 | Linear | Y | -- | 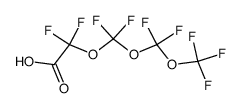 |
| PFO4DA* | PFAA | PFECA | Perfluoro-3,5,7,9-butaoxadecanoic acid | 39492-90-5 | C_6_HF_11_O_6_ | Per | 4 | Linear | Y | -- | 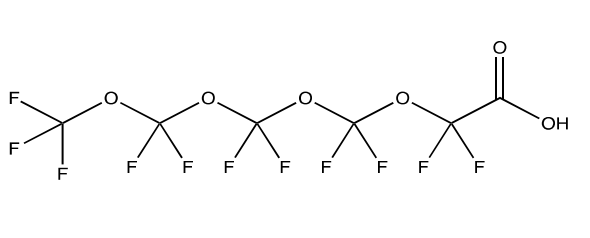 |
| PFO5DoDA  (TAFN4)* | PFAA | PFECA | Perfluoro  (3,5,7,9,11 pentaoxadodecanoic) acid | 39492-91-6 | C_7_HF_13_O_7_ | Per | 5 | Linear | Y | -- | 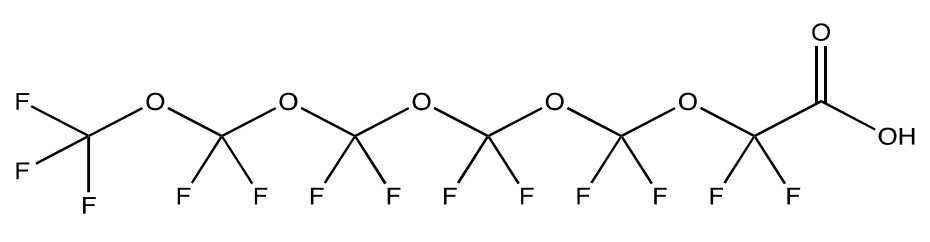 |
| Hydro-EVE Acid* | PFAA | PFECA | Perfluoroethoxsypropanoic acid | 773804-62-9 | C_8_H_2_F_14_O_4_ | Per | 2 | Branched | Y | -- | 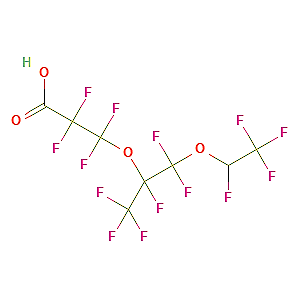 |
| NVHOS* | PFAA | PFESA | Perfluoroethoxysulfonic acid | 1132933-86-8 | C_4_H_2_F_8_O_4_S | Per | 1 | Linear | -- | Y | 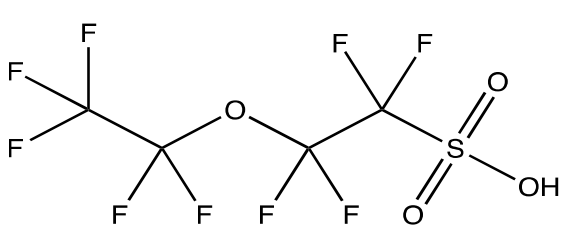 |
| Nafion Byproduct 1* | PFAA | PFESA | Perfluoro-3,6-dioxa-4-methyl-7-octene-1-sulfonic acid | 29311-67-9 | C_7_HF_13_O_5_S | Per | 2 | Branched | -- | Y | 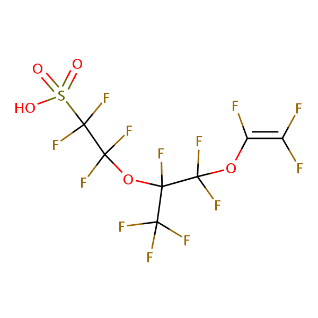 |
| Nafion  Byproduct 2* | PFAA | PFESA | 7H-Perfluoro-4-methyl-3,6-dioxaoctanesulfonic acid | 749836-20-2 | C_7_H_2_F_14_O_5_S | Poly | 2 | Branched | -- | Y | 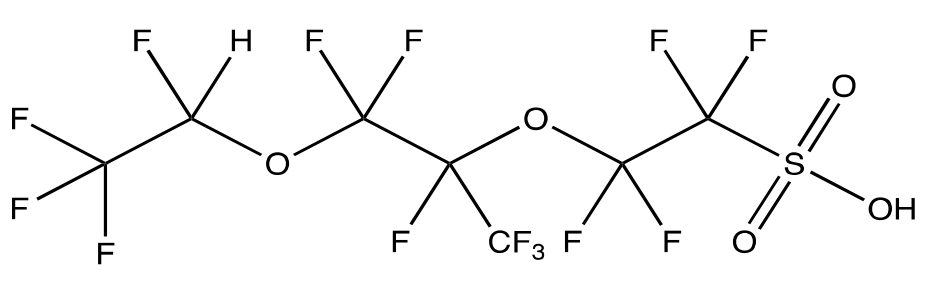 |
| R-PSDA (Formerly Nafion Byproduct 4)* | PFAA | PFESA | Perfluoro-4-(2-sulfoethoxy)pentanoic acid | 2416366-18-0 | C_7_H_2_F_12_O_6_S | Per | 1 | Branched | Y | -- | 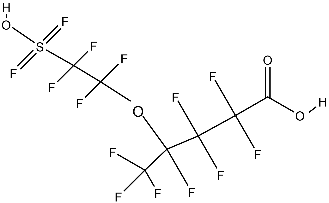 |
| 6:2 FTS | FTSA |  | 6:2 fluorotelomersulfonic acid | 27619-97-2 | C_8_H_5_F_13_O_3_S | Poly | 0 | Linear | -- | Y | 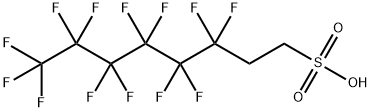 |

a: Carboxylic acid functional group; b: Sulfonic acid functional group. Perfluoroalkyl acid (PFAA); Perfluorocarboxylic acid (PFCA); Perfluorosulfonic acid (PFSA); Perfluoroalkane sulfonamides (FASA); Perfluoroalkane sulfonamidoacetic acids (FASAA); Perfluoroether carboxylic acids (PFECA); Perfluoroether sulfonic acids (PFESA); Fluorotelomer sulfonic acid (FTSA)

**Table S2. Contained in Data File 1.**

| **Table S3. Detected PFAS Concentrations** | | | | | | | | | | | | | | | | | | |
| --- | --- | --- | --- | --- | --- | --- | --- | --- | --- | --- | --- | --- | --- | --- | --- | --- | --- | --- |
| **Location** | **N =** | **Sex (F/M)** |  | **∑ PFAS** | **PFBA** | **PFHxA** | **PFOA** | **PFNA** | **PFDA** | **PFHxS** | **PFOS** | **PMPA** | **HFPODA** | **PFO4DA** | **PFO5DoDA** | **NVHOS** | **Nafion BP2** | **6:2 FTS** |
| Cape Fear River | 48 | 19/29 | Freq (%) | 100 | 83 | 94 | 85 | 98 | 81 | 90 | 100 | 13 | 27 | 46 | 69 | 25 | 79 | 17 |
|  |  |  | Mean (ng/ml) | 166 | 2.04 | 3.19 | 0.38 | 0.85 | 18.47 | 2.16 | 133 | 0.08 | 1.34 | 0.06 | 0.37 | 0.13 | 0.75 | 3.95 |
|  |  |  | Median  (ng/ml) | 27.90 | 1.31 | 0.28 | 0.26 | 0.81 | 3.83 | 0.66 | 12.07 | <LOD | <LOD | <LOD | 0.15 | <LOD | 0.26 | <LOD |
|  |  |  | Min (ng/ml) | 1.78 | ND | ND | ND | ND | ND | ND | 0.65 | ND | ND | ND | ND | ND | ND | ND |
|  |  |  | Max (ng/ml) | 4194 | 26.4 | 44.4 | 1.29 | 8.45 | 466 | 27.2 | 4061 | 1.46 | 15.0 | 0.19 | 1.55 | 0.72 | 7.75 | 95.2 |
| Lake Waccamaw | 27 | 7/20 | Freq (%) | 100 | 37 | 69 | 59 | 89 | 85 | 74 | 100 | 15 | 0 | 22 | 22 | 0 | 4 | 0 |
|  |  |  | Mean (ng/ml) | 10.2 | 0.06 | 0.03 | 0.21 | 0.36 | 1.51 | 0.26 | 7.67 | 0.02 | ND | 0.02 | 0.03 | ND | 0.01 | ND |
|  |  |  | Median  (ng/ml) | 7.66 | <LOD | <LOD | 0.08 | 0.24 | 0.84 | 0.09 | 5.62 | <LOD | ND | <LOD | <LOD | ND | <LOD | ND |
|  |  |  | Min (ng/ml) | 1.15 | ND | ND | ND | ND | ND | ND | 0.25 | ND | - | ND | ND | - | ND | - |
|  |  |  | Max (ng/ml) | 40.5 | 0.26 | 0.08 | 2.82 | 0.91 | 5.00 | 3.35 | 36.9 | 0.30 | - | 0.09 | 0.14 | - | 0.12 | - |

F, female; M, male, Freq, detection frequency; LOD, limit of detection; Min, minimum concentration; Max, maximum concentration; ND, not detected.

**Table S4. SRM 1957 Analysis**

|  | **SRM 1957 (ng/mL)** | | **Certified Value (ng/mL)** | | **% Difference** | |
| --- | --- | --- | --- | --- | --- | --- |
| ***PFOA*** | | 4.812 ± 0.057 | | 5 ± 0.440 | | 3.76 |
| ***PFNA*** | | 0.852 ± 0.018 | | 0.878 ± 0.077 | | 2.96 |
| ***PFDA*** | | 0.328 ± 0.004 | | 0.39 ± 0.12 | | 15.89 |
| ***PFOS*** | | 21.643 ± 0.341 | | 21.1 ± 1.3 | | 2.57 |
| ***PFHxS*** | | 3.652 ± 0.041 | | 4 ± 0.830 | | 8.70 |

NIST Standard Reference Material 1957, ran in triplicate within each batch of samples (n = 6).

| **Table S5. Alligator gene list and results of qRT-PCR mRNA analysis** | | | | | | | |
| --- | --- | --- | --- | --- | --- | --- | --- |
| **Gene Name** | **Gene Symbol** | **RefSeq IDs** | **Possible Product(s)** | **Product Length (bp)** | **Amplification Efficiencies**  **(%; Slope; R^2^)** | **Fold Change (Mean ± SEM)** | **p-value** |
| C-X-C motif chemokine ligand 10 | *Cxcl10* | [XM_006270817.2](https://www.ncbi.nlm.nih.gov/entrez/viewer.fcgi?db=nucleotide&id=1113785784) | C-X-C motif chemokine 10 (LOC102569407) | 117 | 90.9; -3.56; 0.99 | 478 ± 241 | .001 |
| Interferon-induced GTP-binding protein Mx1 | *Mx1* | [XM_019476368.1](https://www.ncbi.nlm.nih.gov/entrez/viewer.fcgi?db=nucleotide&id=1113794399) [XM_019476367.1](https://www.ncbi.nlm.nih.gov/entrez/viewer.fcgi?db=nucleotide&id=1113794397)  [XM_019476366.1](https://www.ncbi.nlm.nih.gov/entrez/viewer.fcgi?db=nucleotide&id=1113794395)  [XM_006274589.3](https://www.ncbi.nlm.nih.gov/entrez/viewer.fcgi?db=nucleotide&id=1113794403)  [XM_019476369.1](https://www.ncbi.nlm.nih.gov/entrez/viewer.fcgi?db=nucleotide&id=1113794401) | interferon-induced GTP-binding protein Mx1 (LOC102568651), transcript variants X1, X2, X3  interferon-induced GTP-binding protein Mx1-like (LOC106736680), transcript variants X1, X2 | 104 | 96.1; -3.42; 0.98 | 405 ± 201 | .009 |
| 2'-5'-oligoadenylate synthase 1 | *Oas1* | [XM_019490991.1](https://www.ncbi.nlm.nih.gov/entrez/viewer.fcgi?db=nucleotide&id=1113764105) | 2'-5'-oligoadenylate synthetase 1, transcript variant X2 | 70 | 88.7; -3.63; 0.97 | 577 ± 162 | .001 |
| Radical S-adenosyl methionine domain-containing protein 2 | *Rsad2* | [XM_019492667.1](https://www.ncbi.nlm.nih.gov/entrez/viewer.fcgi?db=nucleotide&id=1113768703) | radical S-adenosyl methionine domain containing 2 | 110 | 94.8; -3.45; 0.99 | 61.1 ± 20.7 | .004 |
| Glyceraldehyde-3-phosphate dehydrogenase | *Gapdh* | [XM_006258364.3](https://www.ncbi.nlm.nih.gov/entrez/viewer.fcgi?db=nucleotide&id=1113794805) | glyceraldehyde-3-phosphate dehydrogenase | 78 | 95.4; -3.44; 0.99 | - | - |

| **Table S6. Primer sequences used for qRT-PCR analysis.** | |
| --- | --- |
| **Primer** | **Primer sequence (5' - 3')** |
| Cxcl10-F_qRT | TCATGCCTCAGCTAATGTAACC |
| Cxcl10-R_qRT | GTCAGTACAAGCTGCTCTCTTT |
| Cxcl10-P_qRT | AGTGATGACCGTGCTGAAACACCA |
| Mx1-F_qRT | CTGGTGAAGGAGTGGGTATTAG |
| Mx1-R_qRT | CACTCTTGTGATCCCTGGTAAG |
| Mx1-P_qRT | CCCTGGAAATCAGCTCCCAAGATGT |
| Oas1-F_qRT | TCAGAAAACGCAGCCTGAAC |
| Oas1-R_qRT | ATGCTTAGCCACAGAGCGAA |
| Oas1-P_qRT | TCAAAGGGAGCAAGGG |
| Rsad2-F_qRT | ACTACAAGTGCGGCTTCTGC |
| Rsad2-R_qRT | AATTTTCTCCATACCTGCCGCC |
| Rsad2-P_qRT | TTCCACACGGCCAAGA |
| Gapdh-FqRT | AAAGTCGGCGTAAACGGATTTG |
| Gapdh-R_qRT | GACCTCAACTTTGCCACTGC |
| Gapdh-P_qRT | TATCGGCCGCCTGGT |
| F, forward primer; R, reverse primer; P, fluorescein (FAM) labeled TaqMan probe | |

| **Table S7. Results of general linear modeling with main effect of site** | | | | | | |
| --- | --- | --- | --- | --- | --- | --- |
| **Multivariate Tests** | | | | | | |
| Effect | F-statistic | Hypothesis df | Error df | Sig | Partial Eta Squared | Observed Power |
| Intercept | 1.51 | 14 | 33 | 0.16 | 0.39 | 0.7 |
| SVL | 0.92 | 14 | 33 | 0.55 | 0.28 | 0.44 |
| Sex | 1.21 | 14 | 33 | 0.31 |  |  |
| Month | 1.8 | 14 | 33 | 0.08 | 0.154769192 | 0.532299255 |
| Year | 1.51 | 14 | 33 | 0.16 | 0.019669329 | 0.155911039 |
| **Location** | **3.24** | **14** | **33** | **0.003** | **0.011165429** | **0.108723566** |
| **Between-Subject Tests** | | | | | | |
| PFAS | Effect | F-statistic | df | Sig. | Partial Eta Squared | Observed Power |
| PFBA | **Corrected Model** | **2.58** | **5** | **0.04** | **0.22** | **0.75** |
|  | Intercept | 1.89 | 1 | 0.18 | 0.04 | 0.27 |
|  | SVL | 0.44 | 1 | 0.51 | 0.01 | 0.10 |
|  | Sex | 2.79 | 1 | 0.10 | 0.06 | 0.37 |
|  | Month | 0.85 | 1 | 0.36 | 0.02 | 0.15 |
|  | Year | 1.89 | 1 | 0.18 | 0.04 | 0.27 |
|  | Location | 3.83 | 1 | 0.06 | 0.08 | 0.48 |
| PMPA | Corrected Model | 0.89 | 5 | 0.50 | 0.09 | 0.29 |
|  | Intercept | 0.26 | 1 | 0.61 | 0.01 | 0.08 |
|  | SVL | 0.71 | 1 | 0.41 | 0.02 | 0.13 |
|  | Sex | 1.85 | 1 | 0.18 | 0.04 | 0.27 |
|  | Month | 0.00 | 1 | 0.97 | 0.00 | 0.05 |
|  | Year | 0.26 | 1 | 0.61 | 0.01 | 0.08 |
|  | Location | 1.97 | 1 | 0.17 | 0.04 | 0.28 |
| **NVHOS** | Corrected Model | 1.99 | 5 | 0.10 | 0.18 | 0.62 |
|  | Intercept | 0.12 | 1 | 0.73 | 0.00 | 0.06 |
|  | SVL | 0.98 | 1 | 0.33 | 0.02 | 0.16 |
|  | Sex | 0.00 | 1 | 0.95 | 0.00 | 0.05 |
|  | Month | 0.13 | 1 | 0.72 | 0.00 | 0.06 |
|  | Year | 0.12 | 1 | 0.73 | 0.00 | 0.06 |
|  | **Location** | **6.97** | **1** | **0.01** | **0.13** | **0.73** |
| PFHxA | Corrected Model | 0.77 | 5 | 0.57 | 0.08 | 0.25 |
|  | Intercept | 0.53 | 1 | 0.47 | 0.01 | 0.11 |
|  | SVL | 0.18 | 1 | 0.67 | 0.00 | 0.07 |
|  | Sex | 0.01 | 1 | 0.93 | 0.00 | 0.05 |
|  | Month | 1.36 | 1 | 0.25 | 0.03 | 0.21 |
|  | Year | 0.53 | 1 | 0.47 | 0.01 | 0.11 |
|  | Location | 0.98 | 1 | 0.33 | 0.02 | 0.16 |
| GenX | Corrected Model | 1.94 | 5 | 0.11 | 0.17 | 0.60 |
|  | Intercept | 0.20 | 1 | 0.66 | 0.00 | 0.07 |
|  | SVL | 3.81 | 1 | 0.06 | 0.08 | 0.48 |
|  | Sex | 0.44 | 1 | 0.51 | 0.01 | 0.10 |
|  | Month | 2.52 | 1 | 0.12 | 0.05 | 0.34 |
|  | Year | 0.20 | 1 | 0.66 | 0.00 | 0.07 |
|  | Location | 2.31 | 1 | 0.14 | 0.05 | 0.32 |
| **PFO4DA** | **Corrected Model** | **3.94** | **5** | **0.00** | **0.30** | **0.92** |
|  | Intercept | 0.06 | 1 | 0.80 | 0.00 | 0.06 |
|  | SVL | 0.36 | 1 | 0.55 | 0.01 | 0.09 |
|  | Sex | 0.48 | 1 | 0.49 | 0.01 | 0.10 |
|  | **Month** | **7.48** | **1** | **0.01** | **0.14** | **0.76** |
|  | Year | 0.06 | 1 | 0.80 | 0.00 | 0.06 |
|  | Location | 3.73 | 1 | 0.06 | 0.07 | 0.47 |
| 6:2-FTS | Corrected Model | 0.62 | 5 | 0.69 | 0.06 | 0.21 |
|  | Intercept | 0.01 | 1 | 0.93 | 0.00 | 0.05 |
|  | SVL | 0.88 | 1 | 0.35 | 0.02 | 0.15 |
|  | Sex | 0.61 | 1 | 0.44 | 0.01 | 0.12 |
|  | Month | 0.31 | 1 | 0.58 | 0.01 | 0.08 |
|  | Year | 0.01 | 1 | 0.93 | 0.00 | 0.05 |
|  | Location | 0.92 | 1 | 0.34 | 0.02 | 0.16 |
| **PFHxS** | **Corrected Model** | **3.35** | **5** | **0.01** | **0.27** | **0.86** |
|  | Intercept | 0.13 | 1 | 0.72 | 0.00 | 0.06 |
|  | SVL | 0.31 | 1 | 0.58 | 0.01 | 0.08 |
|  | Sex | 0.38 | 1 | 0.54 | 0.01 | 0.09 |
|  | **Month** | **6.42** | **1** | **0.01** | **0.12** | **0.70** |
|  | Year | 0.13 | 1 | 0.72 | 0.00 | 0.06 |
|  | **Location** | **7.37** | **1** | **0.01** | **0.14** | **0.76** |
| **PFOA** | Corrected Model | 1.68 | 5 | 0.16 | 0.15 | 0.53 |
|  | Intercept | 0.92 | 1 | 0.34 | 0.02 | 0.16 |
|  | SVL | 0.52 | 1 | 0.47 | 0.01 | 0.11 |
|  | Sex | 0.96 | 1 | 0.33 | 0.02 | 0.16 |
|  | Month | 2.15 | 1 | 0.15 | 0.04 | 0.30 |
|  | Year | 0.92 | 1 | 0.34 | 0.02 | 0.16 |
|  | **Location** | **5.33** | **1** | **0.03** | **0.10** | **0.62** |
| **Nafion_bp2** | Corrected Model | 4.31 | 5 | 0.00 | 0.32 | 0.94 |
|  | Intercept | 1.68 | 1 | 0.20 | 0.04 | 0.24 |
|  | SVL | 0.07 | 1 | 0.79 | 0.00 | 0.06 |
|  | Sex | 0.21 | 1 | 0.65 | 0.00 | 0.07 |
|  | Month | 1.75 | 1 | 0.19 | 0.04 | 0.25 |
|  | Year | 1.68 | 1 | 0.20 | 0.04 | 0.25 |
|  | **Location** | **17.75** | **1** | **0.00** | **0.28** | **0.98** |
| **PFO5DoDA** | **Corrected Model** | **4.35** | **5** | **0.00** | **0.32** | **0.94** |
|  | Intercept | 0.70 | 1 | 0.41 | 0.01 | 0.13 |
|  | SVL | 3.84 | 1 | 0.06 | 0.08 | 0.48 |
|  | Sex | 0.16 | 1 | 0.69 | 0.00 | 0.07 |
|  | Month | 0.23 | 1 | 0.63 | 0.00 | 0.08 |
|  | Year | 0.70 | 1 | 0.41 | 0.02 | 0.13 |
|  | **Location** | **9.17** | **1** | **0.00** | **0.17** | **0.84** |
| **PFNA** | **Corrected Model** | **4.01** | **5** | **0.00** | **0.30** | **0.92** |
|  | Intercept | 3.97 | 1 | 0.05 | 0.08 | 0.50 |
|  | SVL | 0.18 | 1 | 0.67 | 0.00 | 0.07 |
|  | Sex | 1.04 | 1 | 0.31 | 0.02 | 0.17 |
|  | Month | 1.04 | 1 | 0.31 | 0.02 | 0.17 |
|  | Year | 3.97 | 1 | 0.05 | 0.08 | 0.50 |
|  | **Location** | **15.48** | **1** | **0.00** | **0.25** | **0.97** |
| **PFDA** | **Corrected Model** | **2.96** | **5** | **0.02** | **0.24** | **0.81** |
|  | Intercept | 1.83 | 1 | 0.18 | 0.04 | 0.26 |
|  | SVL | 0.00 | 1 | 1.00 | 0.00 | 0.05 |
|  | Sex | 0.57 | 1 | 0.45 | 0.01 | 0.11 |
|  | Month | 2.44 | 1 | 0.13 | 0.05 | 0.33 |
|  | Year | 1.83 | 1 | 0.18 | 0.04 | 0.26 |
|  | **Location** | **10.30** | **1** | **0.00** | **0.18** | **0.88** |
| **PFOS** | Corrected Model | 2.09 | 5 | 0.08 | 0.19 | 0.64 |
|  | Intercept | 2.75 | 1 | 0.10 | 0.06 | 0.37 |
|  | SVL | 0.01 | 1 | 0.94 | 0.00 | 0.05 |
|  | Sex | 0.00 | 1 | 0.97 | 0.00 | 0.05 |
|  | Month | 2.06 | 1 | 0.16 | 0.04 | 0.29 |
|  | Year | 2.75 | 1 | 0.10 | 0.06 | 0.37 |
|  | **Location** | **5.82** | **1** | **0.02** | **0.11** | **0.66** |
| GLM with main effect of location (LW or CFR) with random effect of SVL, sex, month, and year. All F-statistic show Roy's Largest Root result of the multivariate analysis of variance. Effects that are significant (p < .05) are bolded. | | | | | | |
